# High genetic diversity in the pelagic deep-sea fauna of the Atacama Trench revealed by environmental DNA

**DOI:** 10.1101/2022.04.14.488404

**Authors:** Salvador Ramírez-Flandes, Carolina E. González, Montserrat Aldunate, Julie Poulain, Patrick Wincker, Ronnie N. Glud, Rubén Escribano, Sophie Arnaud Haond, Osvaldo Ulloa

## Abstract

A current paradigm in marine biodiversity states that faunal richness decreases with depth. However, the deep-ocean ecosystem has been significantly under-sampled, hindering a complete view of its biodiversity. This situation is accentuated in ultra-deep waters, where the remote and extreme conditions unfit the traditional sampling methods. Using environmental DNA, we assessed the pelagic metazoan diversity of the Atacama Trench from the high-productive near-surface level down to ~8000 m depth. Our results show that waters deeper than 4000 m contributed up to 50% of the overall genetic diversity. These findings contrast with similar observations in the less-productive Kermadec Trench, where the diversity in deep waters was lower than in shallower waters. Moreover, both deep pelagic ecosystems exhibited some unknown phylogenetic clades within the dominant taxonomic groups: hydrozoans and copepods. The deep-ocean biota may thus contribute to global biodiversity far more than hitherto suggested, especially in zones influenced by high primary production. Our results underline the need for increased effort to study these remote ecosystems and improve our understanding of their contribution to the ecology and biogeochemistry of the deep-sea pelagic and benthic realms.

## Introduction

Knowledge of the global magnitude, geographic distribution, and biodiversity loss is essential to ecology and conservation biology. This need has dramatically enhanced the number of biogeographic investigations in the past decades (Gaston, 2000). Current estimates suggest that about two-thirds of the species on Earth have been described so far (Costello and Chaudhary, 2017). The number of species has been estimated at ~0.3 million in the ocean and ~1.7 million on land (including fresh-water ecosystems) (Costello and Chaudhary, 2017), yet the forecast of the remaining undescribed diversity is highly variable, particularly in the marine realm (Appeltans et al., 2012). These estimates are partially based on the fact that only ~16% of currently named species are marine (Grosberg et al., 2012), challenging the classical biogeographic model that relates habitat size to species richness (Costello and Chaudhary, 2017). However, less than 1% of the deep ocean has been explored (Higgs and Attrill, 2015), due primarily to the logistic and technological difficulties involved in sampling the vast and high-hydrostatic-pressure abyssal (4,000—6,000 m) and hadal (6,000—11,000 m) realms (Smith et al., 2009). Moreover, the deep sea is characterized by low biomass production and import, and it has long been expected to be scarcely populated with a handful of scavengers thriving in a cold and biomass-poor world (Ramirez-Llodra et al., 2010). Consequently, our knowledge of the faunal diversity below 1 km depth—almost 80% of the biosphere (Childress, 1995)—is elusive. Indeed, as our comprehension of the latitudinal gradient in marine biodiversity is biased by sampling effort (Chaudhary et al., 2016), that of the vertical distribution of marine biodiversity is also similarly biased. The deep-sea diversity has therefore been suggested to equal or even surpass that present in shallow waters (Costello and Chaudhary, 2017; Valentine and Jablonski, 2015), despite the hostile conditions of the deep sea, which is cold, dark, extremely diluted, and contains limited food sources (Jamieson et al., 2010; Priede and Froese, 2013). Conversely, other studies suggest that this comparatively unbalanced picture of species richness reflects an ecological truth rather than unbalanced sampling efforts—unless many rare and endemic species would be discovered in the deep sea or, for example, if marine parasites were far more diverse than presently documented (Costello and Chaudhary, 2017). Moreover, hadal trenches sustain elevated benthic biological activity at extreme hydrostatic pressure due to material focusing (Glud et al., 2013), but their pelagic faunal realm remains unexplored.

In the last decade, marine ecologists have started to assess marine metazoan biodiversity using methods based on the analysis of environmental DNA (eDNA) (Deiner et al., 2017) to complement the traditional methods based on the sampling and study of entire organisms. This molecular method is based on the fact that organisms leave traces of their genetic material in the environment through shedding and waste depositions (Taberlet et al., 2012). The use of eDNA allows the detection of high levels of biodiversity with a reasonable amount of starting material, thus offering a way to circumvent the problem of sampling the deep sea. Since DNA degradation in the water column typically occurs within only a few days (Andruszkiewicz Allan et al., 2021; Harrison et al., 2019; Thomsen et al., 2012), this technique can allow estimates of the biodiversity present at the sampling time or, at least, within a relatively short time window. Thus, eDNA-based tools have emerged as a promising alternative for rapid, sensitive, and non-invasive assessment of biodiversity (Harrison et al., 2019). However, to date, marine eDNA metazoan biodiversity studies have mainly focused on the epipelagic zone (< 200 m depth) and the ocean seafloor sediments (Cristescu and Hebert, 2018; Sinniger et al., 2016), leaving unexplored the majority of the water column—from the mesopelagic zone (200—1000 m) down to the abyssal and hadal zones (4,000— 11,000 m). Recently, the eDNA method has been applied at the basin scale to assess the pelagic eukaryotic diversity, including the very abundant protists, down to 5200 m, and compare it with that in the abyssal and bathyal surficial sediments (Cordier et al., 2022). The composition of pelagic metazoan communities in the bathypelagic or abyssal, let alone the hadal, realm thus remains poorly known despite their potential importance for the stability of ecosystems, as for models for climate change and conservation policies. Here we used eDNA metabarcoding of two genetic markers, nuclear 18S rRNA and mitochondrial cytochrome c oxidase I genes, henceforth, 18S and COI, to assess the metazoan biodiversity in the water column of the Atacama Trench—from 85 m to more than 8000 m depth—and in the abyssal and hadal waters of the Kermadec Trench (Atacama and Kermadec, hereinafter; Fig. 1 and Table S1).

**Figure 1.**
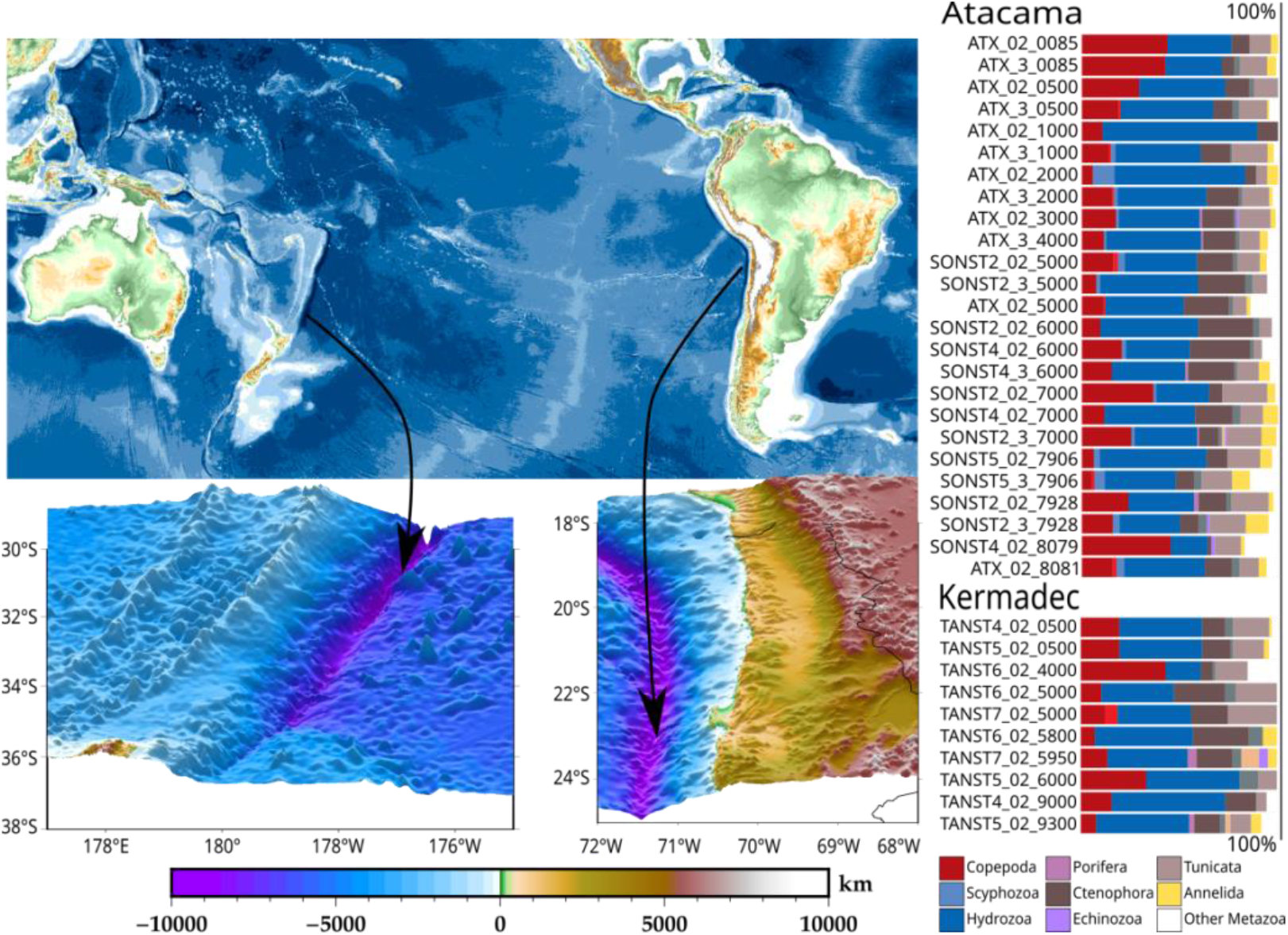
Geographic setting of the sampling of eDNA data in the Kermadec and Atacama Trenches (left) and taxonomic diversity distribution (presence/absence) of the Amplicon Sequence Variants (ASVs) of the 18S rRNA gene metabarcoding (right). The names of the samples are split into three parts, encoding the name of the cruise/station, the water filter size range (“02” for the range 0.2-3 μm, and “3” for the range 3-20 μm), and the depth (meters). Detailed information on the samples can be found in Table S1.

## Results

The analysis of the combined data (12,093,326 and 11,551,435 18S and COI metabarcoding sequences, respectively) resulted in 497 metazoan 18S Amplicon Sequence Variants (ASVs) and 307 COI translated ASVs (tASVs). The taxonomic diversity of these sequences was mainly associated with the cnidarian class Hydrozoa and the crustacean subclass Copepoda throughout the water column, while tunicates, annelids, scyphozoans, placozoans, and sponges were also prominent in some samples (Figs. 1 and S1; Table S2).

The overall genetic richness distribution of 18S ASVs in the Atacama Trench was slightly higher in the abysso-hadal waters (>4000 m) than in shallower waters and markedly higher compared to the abysso-hadal waters of the Kermadec Trench (Figs. S4 and S5). In Atacama, the metazoan diversity of ASVs was distributed among ~10 metazoan phyla. The majority of these ASVs were found in waters > 4000 m, according to the 18S and COI metabarcoding data (Tables 1 and S2, respectively). An evaluation of the contribution of each depth level to the overall water-column genetic richness revealed that the abysso-hadal waters of the Atacama Trench region harbor ~50% of the total amount of detected ASVs, suggesting a high level of endemism (Fig. 2A and 2B). In contrast, only <20% of the ASVs found in the abysso-hadal samples of Kermadec were not observed at 500 m. Moreover, the percentages of exclusive ASVs per depth followed a similar pattern (Fig. 2C), and consequently, the genetic diversity was observed to be more equally distributed with depth in Atacama than in Kermadec (Fig. 2d).

**Figure 2.**
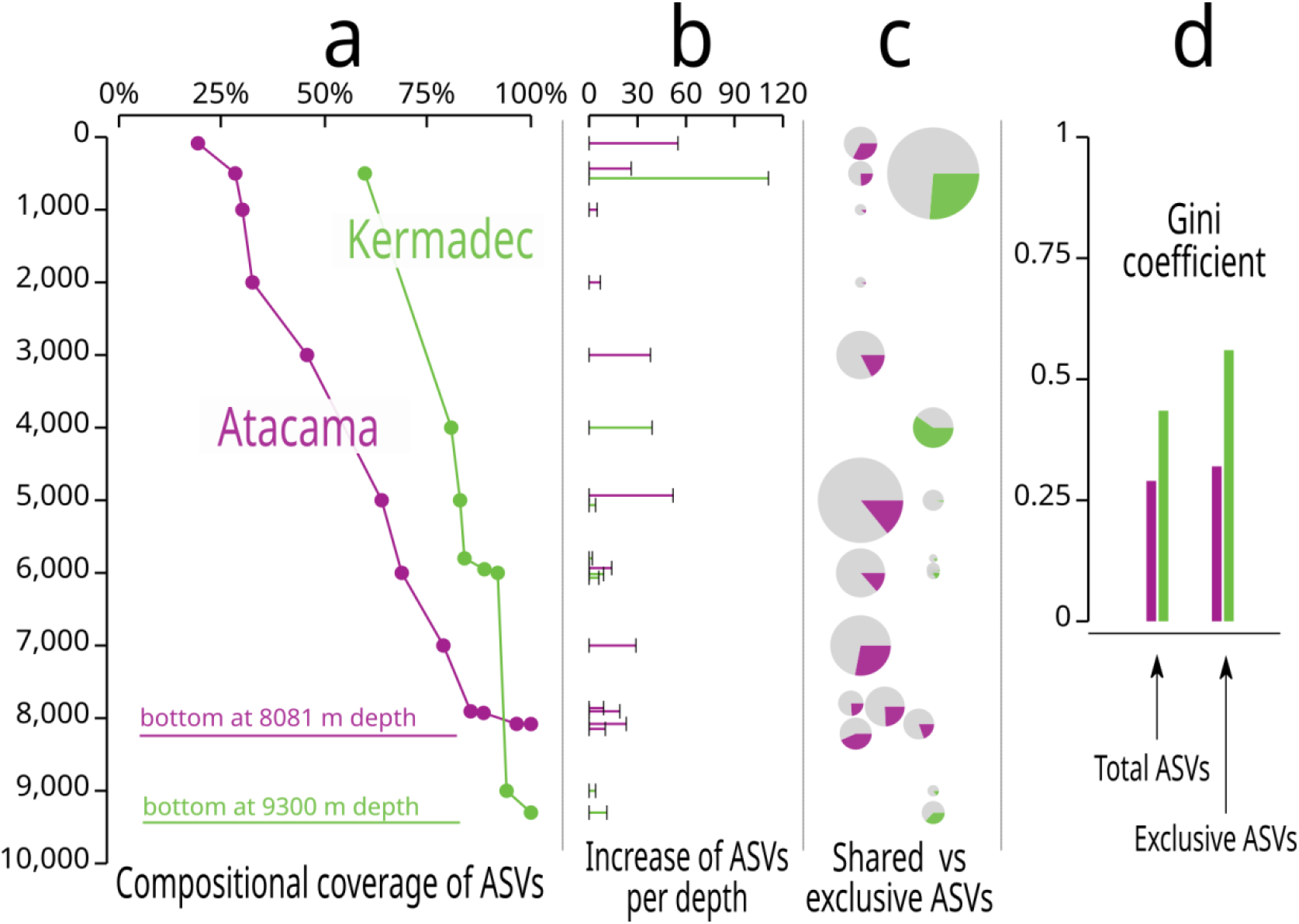
**a)** Compositional coverage of total metazoan ASVs per sample (excluding those obtained with the filter 3-20 μm for Atacama as this filter was not used in Kermadec) shows that the faunal genetic diversity is more uniformly distributed in the water column of Atacama than in Kermadec. For example, at 500 m, the Kermadec system already covered 50% of its metazoan diversity, while in Atacama this was achieved only at abysso-hadal depths. **b)** The per-sample contribution of ASVs in the water columns of Atacama and Kermadec also shows a different pattern, in which samples from the abysso-hadal pelagic system contribute more to the metazoan biodiversity in Atacama than in Kermadec. In this case, the bar lengths represent the number of ASVs of the 18S genetic marker. **c)** Proportion of shared ASVs vs. exclusive number of ASVs per sample. The circles’ sizes represent the total number of ASVs per sample, with the gray slice representing the shared ASVs, and the colored slices depicting the exclusive ASVs (those only present in the corresponding sample.) **d)** The Gini coefficient represents degree of inequality in the distribution of the richness of ASVs between the samples from Atacama and Kermadec. The total and exclusive richness of ASVs was more equally distributed in Atacama than in Kermadec, as this coefficient expresses greater equality with values closer to zero.

**Table 1.** Counts of 18S ASVs per metazoan phyla obtained in Atacama above, below, and above-and-below 4000 m (fourth, fifth, and sixth column, respectively). The second and third columns show the biomes documented for each corresponding phylum. A taxonomic affiliation of the ASVs associated with all the groups can be found in Table S3 for Atacama and Table S4 for Kermadec. Table S2 displays similar information for the COI tASVs.

| Phylum | Aquatic Biomes | General Biomes | <4000m | >4000m | Ubiq. | Total |
| --- | --- | --- | --- | --- | --- | --- |
| **Annelida (mostly polychaetes)** | Pelagic/Benthic | Marine/Freshwater/Terrestrial | 1 | 21 | 7 | 29 |
| **Arthropoda (mostly copepods)** | Pelagic/Benthic | Marine/Freshwater/Terrestrial | 38 | 65 | 42 | 145 |
| **Brachiopoda** | Benthic | Marine | 0 | 0 | 0 | 0 |
| **Bryozoa** | Pelagic/Benthic | Marine/Freshwater | 0 | 0 | 0 | 0 |
| **Chaetognatha** | Pelagic/Benthic | Marine | 0 | 13 | 4 | 17 |
| **Chordata (mostly tunicates)** | Pelagic/Benthic | Marine/Freshwater/Terrestrial | 8 | 18 | 40 | 66 |
| **Cnidaria (mostly hydrozoans)** | Pelagic/Benthic | Marine/Freshwater | 21 | 41 | 66 | 128 |
| **Ctenophora** | Pelagic/Benthic | Marine/Freshwater | 0 | 3 | 1 | 4 |
| **Cycliophora** | Benthic | Marine | 0 | 0 | 0 | 0 |
| **Echinodermata** | Benthic | Marine | 1 | 4 | 0 | 5 |
| **Entoprocta** | Pelagic/Benthic | Marine/Freshwater | 0 | 0 | 0 | 0 |
| **Gastrotricha** | Benthic | Marine/Freshwater | 0 | 0 | 0 | 0 |
| **Gnathostomulida** | Benthic | Marine | 0 | 0 | 0 | 0 |
| **Hemichordata** | Pelagic/Benthic | Marine | 0 | 3 | 0 | 3 |
| **Micrognathozoa** | Pelagic/Benthic | Freshwater | 0 | 0 | 0 | 0 |
| **Mollusca** | Pelagic/Benthic | Marine/Freshwater/Terrestrial | 0 | 5 | 1 | 6 |
| **Nematoda** | Benthic | Marine/Freshwater/Terrestrial | 0 | 0 | 0 | 0 |
| **Nemertea** | Benthic | Marine/Freshwater/Terrestrial | 0 | 0 | 0 | 0 |
| **Onychophora** | NA | Terrestrial | 0 | 0 | 0 | 0 |
| **Phronida** | Benthic | Marine | 0 | 0 | 0 | 0 |
| **Placozoa** | Pelagic/Benthic | Marine | 0 | 0 | 0 | 0 |
| **Platyhelminthes** | Pelagic/Benthic | Marine | 0 | 0 | 0 | 0 |
| **Porifera (sponges)** | Pelagic/Benthic | Marine/Freshwater | 0 | 3 | 1 | 4 |
| **Priapulida** | Benthic | Marine | 0 | 0 | 0 | 0 |
| **Syndermata** | Pelagic/Benthic | Marine/Freshwater | 0 | 0 | 0 | 0 |
| **Tardigrada** | Pelagic/Benthic | Marine/Freshwater/Terrestrial | 0 | 0 | 0 | 0 |
| **Xenacoelomorpha** | Pelagic/Benthic | Marine | 0 | 0 | 0 | 0 |
| **Total** |  |  | **69** | **176** | **162** | **407** |

The phylogenetic relationships of the 18S ASVs associated with the taxonomic groups with the highest genetic diversity—namely, Hydrozoa and Copepoda (Figs. 1, S1, S4, and S5)—with the closest related sequences present in Genbank, revealed multiple well-supported (>95% of probability) clades exclusively composed of ASVs observed in this study (Figs. 3, S6 and S7). One of these clades (Hyd_Isedis1, Fig. 3a) contained ASVs exclusively detected in Atacama that could not be placed within any recognized hydrozoan order, suggesting that it might represent a novel *incertae sedis* clade within the Hydroidolina subclass. Other such putative novel hydrozoan clades exclusively comprising ASVs recovered from Atacama were found within Siphonophorae (Hyd_siph1 and Hyd_siph2, Fig. S6a), Rhopalonematidae (Hyd_rhop1, Fig. S6b), and Geryonidae (Hyd_gery1, Fig. S6c). A putative novel hydrozoan clade exclusively composed of ASVs from Kermadec was found within Clytiidae (Hyd_clyt1, Fig. S6d). Also, novel copepod clades with an exclusive composition of ASVs recovered from Atacama were found within Oncaeidae (Cop_onca1 and Cop_onca2, Fig. S7a), Clausocalanidae (Cop_clau1, Fig. S7b) and Calanidae (Cop_cala1 and Cop_cala2, Fig. S7c).

**Figure 3.**
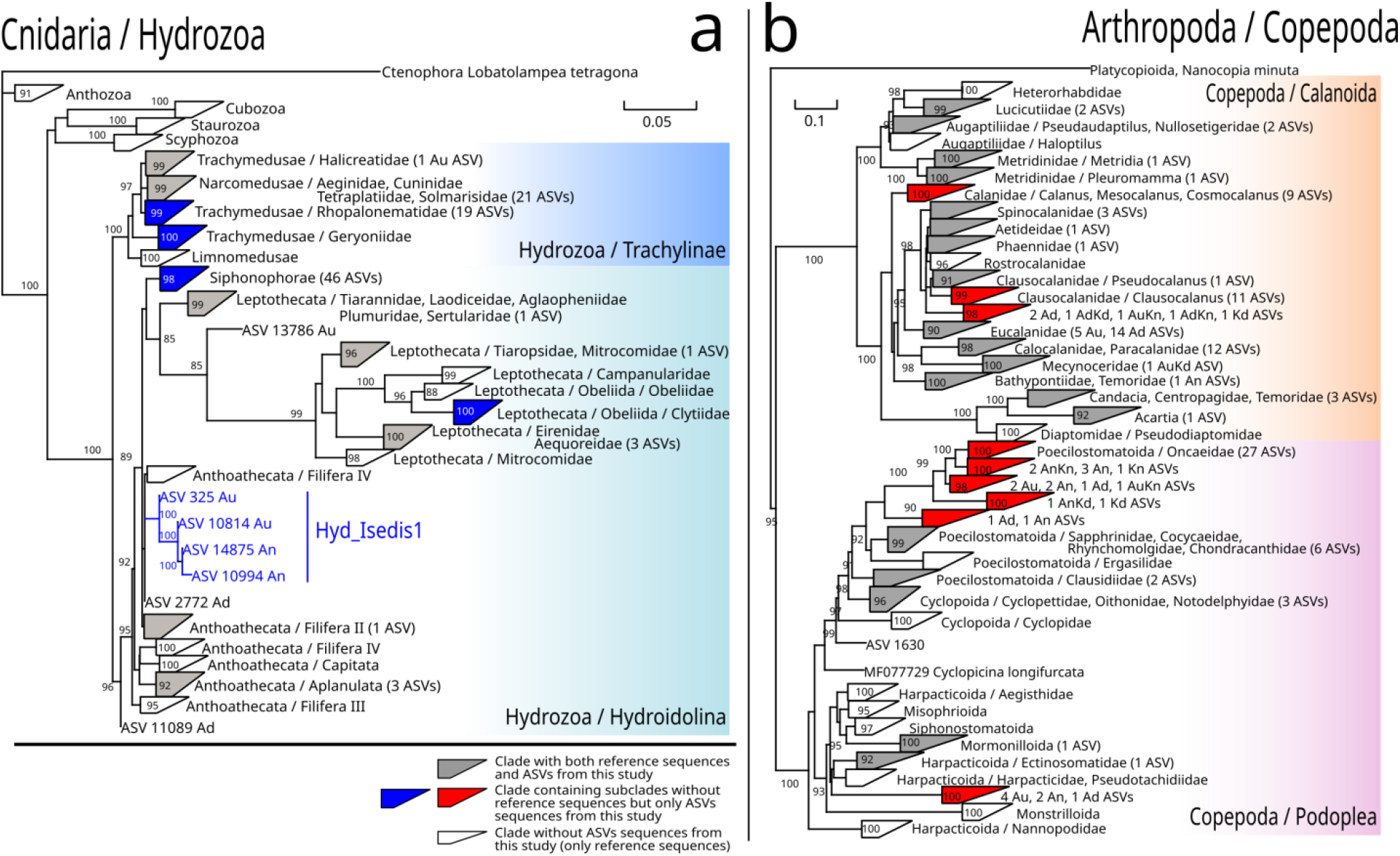
Phylogenetic trees of the 18S ASVs associated with Hydrozoa (a) and Copepoda (b). The whole set of reference sequences utilized for these trees can be found in Table S6 and Table S7 for Hydrozoa and Copepoda, respectively. The colored clades in which the ASVs generated exclusive groups are expanded in Figs. S6 and S7, for Hydrozoa and Copepoda, respectively.

The diversity of the 18S ASVs within Hydrozoa and Copepoda was also notably higher in Atacama than in Kermadec (Figs. S2 and S3). Regarding Hydrozoa, although the taxonomic diversity and abundance of eDNA sequences of siphonophores (pelagic colonial hydrozoans) was predominantly observed in Atacama and Kermadec, the overall taxonomic diversity at the family level was richer in Atacama than in Kermadec (Figs. S2a and S3a). Moreover, a significant proportion of eDNA sequences associated with calycophoran siphonophores were replaced by physonect siphonophores at hadal depths (Fig. S2a and S3a).

Concerning copepods, calanoids of the families Paracalanidae, Calocalanidae, and Eucalanidae prevailed over other calanoid and podoplean groups at most depths (Fig. S2b). In Kermadec, the most prominent calanoid groups were Lucicutiidae, Clausocalanidae, and Mecynoceridae, along with a strong dominance of harpacticoids affiliated with Pseudotachidiidae, persistently observed in the two deepest samples (Figs. S2b and S3b). Pseudotachidiids have been reported in several abyssal ecosystems (Kim et al., 2019) and geographically diverse deep-sea sediments (Sinniger et al., 2016), but our study did not detect them in Atacama. Notably, the eDNA signature of the copepod community in Atacama at 500 m depth was remarkably different from the signature of those in any other sample of this work. This sample came from the lower limit of the Anoxic Marine Zone (AMZ) situated in the eastern South Pacific Ocean (Ulloa et al., 2012)—an oceanographic feature absent in the Kermadec Trench region. This eDNA AMZ signature pointed to metridinids and oncaeids, whose adaptations to low-oxygen conditions—most likely to avoid predation from organisms with higher oxygen demands— have been described in other hypoxic marine ecosystems (Böttger-Schnack et al., 2008; Teuber et al., 2013).

Ctenophores (comb jellies) and Tunicates were also conspicuous in Atacama and Kermadec (Fig. 1). The phylogenetic analysis associated with ctenophores did not form well-supported clades of ASVs, as it did in the case of tunicates. This analysis classified the ASVs of tunicates within the Appendicularia and Thaliacea (pelagic tunicates) classes and none within the Ascidacea (benthic tunicates) class. In the Appendicularia class, two clades exclusively composed of ASVs from this study were grouped (Tun_App1 and Tun_App2), and several ASVs remained unattached to any known family within this class (Fig. S8). The rest of the taxonomic groups mainly included, among others, polychaetes in Annelida; decapods and euphausiids in Crustacea; saggitoideans in Chaethognatha; fishes from the families Atherinidae, Carangidae, Gobiidae, and Sebastidae within chordates from Actinopteri; and murex snails, tulip snails, and sea butterflies in Mollusca (Tables S3 and S4).

## Discussion

The coarse-level taxonomic pattern of the metazoan diversity detected in the Atacama and Kermadec trenches (Figs. 1 and S1) is consistent with those reported for other related pelagic ecosystems, which were based on studies conducted on entire organisms collected with net tows down to 4000 m depth (de Puelles et al., 2019; Kosobokova and Hirche, 2000; Vereshchaka et al., 2017) or inferred from molecular taxonomic markers of epi-mesopelagic (< 1000 m) marine snow (Lundgreen et al., 2019). However, the distribution of the genetic richness of the 18S ASVs in the Atacama Trench was slightly higher in the abysso-hadal waters (>4000 m) compared to shallower waters and markedly higher compared to the abysso-hadal waters of the Kermadec Trench (Figs. S4 and S5). This result contrast with results reported down to mesopelagic depths in other regions using traditional methods (Costello and Chaudhary, 2017; Jamieson et al., 2010), which generally have shown that the faunal richness decreases with ocean depth—despite never having scrutinized metazoan diversity in waters deeper than 5000 m before.

The abysso-hadal realm represents a significant fraction of the biosphere, yet it is the least-explored environment on Earth (Robison, 2009). This biome is ecologically relevant as it sustains a crucial pelagic food web with a significant role in energy export to the ocean’s sediments (Steinberg and Landry, 2017). The traditional techniques for sampling marine metazoan organisms require specialized sampling gear and lengthy, time-consuming sorting and taxonomic identifications by scarce experts. They are thus, by themselves, insufficient to cover the vast ocean. Recently, new eDNA-based methods have been used in deep ocean sediments and pelagic waters studies. These methods complement the traditional methods as they provide a rapid and more extensive way to assess biodiversity in these vast and remote ecosystems.

To this end, our results highlight two main points. First, hydrozoans and copepods dominate the metazoan diversity throughout these ultra-deep water columns. The predominance of hydrozoan siphonophores in the hadopelagic waters of the Atacama Trench was an unexpected result, as these animals were not, thus far, considered as characteristic of these ecosystems (Jamieson, 2015). This result should be interpreted with care, though, as mass jelly die-offs—called jelly-falls— significantly contribute to the pelagic particulate matter that ends on the seabed (Lebrato et al., 2012; Luo et al., 2020). However, the persistence of this signature across sites spanning several kilometers and the distinct type of siphonophores found in hadal (as compared to abyssal) waters in Atacama (Figs. S2a and S3a) make it more likely that these eDNA signatures come from indigenous organisms. Moreover, a significant proportion of eDNA sequences associated with calycophoran siphonophores were replaced by physonect siphonophores at hadal depths (Fig. S2a and S3a). Unlike calycophores, physonect siphonophores possess pneumatophores, which provide their colonies with improved motor skills (Castellani and Edwards, 2017). Other prominent hydrozoans were trachymedusae affiliated with the Rhopalonematidae and Geryoniidae families, whose eDNA sequences generated multiple well-supported phylogenetic clades of ASVs (Fig. 3a), suggesting the presence of several undescribed high-rank taxonomic groups.

Secondly, abyssal and hadal waters in systems like Atacama can contribute to a significant proportion of the overall metazoan genetic diversity in the water column, suggesting the existence of high diversity and endemism below 4000 m depth. The main characteristic of this ecosystem is its high primary production in near-surface waters (Daneri et al., 2000), which might promote a habitat complexity capable of contributing to the development of low-oxygen zones (Ulloa et al., 2012) and making the whole water-column eutrophic (Danovaro et al., 2002), thus promoting biodiversity (Clarke and Gaston, 2006; Grosberg et al., 2012; Vereshchaka et al., 2017).

The models for climate change predictions need to incorporate the diversity and biomass of all the trophic levels in the water column, yet there is a large gap in knowledge about the contribution of metazoans to abyssal and hadal pelagic environments. Diversity promotes stability in ecosystems (Ives and Carpenter, 2007), and thus it represents necessary information that should guide the assessment of the impact of environmental perturbations. This information would improve the output of the ecological models and, consequently, conservation policies. For instance, a recent study suggests that jellies biomass should be included to improve the accuracy of biogeochemical models for carbon estimations (Luo et al., 2020). Moreover, a biodiversity loss in the deep ocean might be associated with an exponential reduction of its ecosystem functions (Danovaro et al., 2008).

The output of any eDNA study depends on the geographical coverage and the primers’ specificity for DNA amplification, as some natural taxonomic groups might have evolved molecular sequences incompatible with the primers. Admittedly, our findings are based on a limited number of hard-to-get samples and a methodology that may still be giving an incomplete view of the actual metazoan diversity present in the environment. Also, the completeness of the available sequence databases severely constrains this view. Nonetheless, our results provide the first view of the diversity and distribution of ultra-deep pelagic fauna and suggest that the deep-ocean biota might contribute to global biodiversity much more than what is calculated in present-day estimates, especially in general zones of increased habitat complexity due to the influence of high primary production. These results also highlight the potential of eDNA-methods to explore marine biodiversity in the three dimensions of the world’s ocean and are consistent with previous sediment observations for the Atacama Trench as a hotspot of faunal diversity (Danovaro et al., 2002). Furthermore, they stress the need for developing and improving standardized approaches to improve our understanding of the ecology of this vast and still underexplored biome.

## Material and Methods

### Collecting samples

Seawater samples were collected during three oceanographic cruises (TAN1711, ATACAMEX and SO261) conducted in the Kermadec and Atacama trenches of the South Pacific Ocean. The TAN1711 with RV *Tangaroa* cruise (from November 24^th^ to December 14^th^, 2017) covered a transect from New Zealand (41°18’S, 174°47’W) to the Northern Kermadec Trench (21°01’S, 175°12’W), involving four oceanographic stations. Both the ATACAMEX cruise with the RV *Cabo de hornos* (from January 28^th^ to February 4^st^, 2018) and SO261 cruise with RV *Sonne* (March 2^nd^ to April 2^nd^, 2018) were carried out in the deepest region of the trench (20°19’S — 24°15’S off northern Chile, Fig. 1, Table S1). Each sample was filtered across a hydrophilic 20-μm nylon membrane (Merck Millipore, Massachusetts, USA) and filtered onto a Sterivex fileter using a peristaltic pump. The membrane filter containing captured eDNA and cellular material from the water column was filled with lysis buffer and stored at −80°C. **DNA extractions** were carried out by Genoscope (Évry, France) using the same protocol as described by Alberti et al. (2017) for Tara Oceans water samples. The protocol is based on the cryogenic grinding of membrane filters, followed by nucleic acid extraction with NucleoSpin RNA kits combined with the NucleoSpin DNA buffer set (Macherey– Nagel, Düren, Germany). **PCR amplification and sequencing:** The 18S-V1V2 metabarcodes were generated using the SSUF04 and SSUR22mod primers. The COI metabarcodes were generated using the mlCOIintF and jgHCO2198 primers. Paired-end sequencing (2×250bp) of the libraries was performed on either the HiSeq 4000 or HiSeq 2500 instruments (Illumina, San Diego, CA, USA). **Sequence analysis:** Paired-end reads were assembled and quality filtered with vsearch v2.17.1, using parameter of maxEE at 0.5, minimum length of 250 bp, and a maximum length of 500 bp. The primers in the resulting metabarcoding sequences were removed with pTrimmer v1.3.4. The sequences lacking the primers were discarded. The sequences from all datasets were then put into a single file, and the unique sequences were determined with vsearch. Chimeras were then removed with the same program, and in the same process, the Amplicon Sequence Variants (ASVs) were created. The 18S ASVs were compared as nucleotide sequences using BLASTN (parameters task dc-megablast, bitscore cutoff of 50) against the SILVA database (138.1) to construct the corresponding taxonomic profiles. The swarm clusters were created with swarm v3.0 with d = 1 and fastidious parameters. The COI ASVs were translated into protein sequences (translated ASV or tASV), using the longest translation of the six open reading frames. These tASVs were aligned against a custom reference sequence database of COI sequences with BLASTP (bitscore cutoff of 50). **Phylogenetic analyses**: The phylogenetic trees of 18S ASVs and references sequences were constructed as follows. First, reference sequences for hydrozoans, copepods, and tunicates were collected from the literature and from respective searches in Genbank, generating separate datasets for each taxonomic group (Tables S5, S6, and S7). These datasets were complemented with the ASVs sequences associated with each taxonomic group according to the previously described procedure (comparison with the SILVA database). Second, these datasets of sequences were then aligned using mafft with parameters for high accuracy. The respective alignments were used to create phylogenetic trees with IQ-TREE2. Further details on these methods and corresponding references can be found in the Supplementary Material.

## Supporting information

Figures and Tables starting with "S".

## Acknowledgments

This work was supported the Chilean National Agency for Research and Development through the Millenium Science Initiative-ANID Program (grant ICN12 019-IMO) and grant Fondecyt 1191360 (to O.U.) and AUB17002 to Wolfgang Schneider. We thank the captains, crews, and scientific personnel of the RV *Sonne* (SO261; ship time provided by BMBF, Germany, awarded to Frank Wenzhoefer, Mathias Zabel, and Ronnie N. Glud), and RV *Tangaroa* (TAN1711; shiptime partly funded by the Coasts & Oceans Centre of New Zealand’s National Institute of Water & Atmospheric Research -NIWA-, awarded to Ashley A. Rowden and Ronnie N. Glud), in the framework of the HADES-ERC Advanced grant “Benthic diagenesis and microbiology of hadal trenches”#669947 awarded to Ronnie N. Glud. Additional support was provided by the Danish National Research Foundation via the Danish Center for Hadal Reserch, grant number DNRF145. This work is part of the “Pourquoi Pas les Abysses?” project funded by Ifremer, and the project eDNAbyss (AP2016 −228) funded by France Génomique (ANR-10-INBS-09) and Genoscope-CEA.

