## Supplementary material for "High genetic diversity in the pelagic deep-sea fauna of the Atacama Trench revealed by environmental DNA": Figures and Tables starting with "S".

### 1. Collection of samples

Seawater samples for eDNA analysis were collected with Niskin bottles during three oceanographic cruises (TAN1711, ATACAMEX and SO261) conducted in the Kermadec and Atacama trenches of the South Pacific Ocean. The TAN1711 cruise with the RV *Tangaroa* (from November 24th to December 14th, 2017) covered a transect from New Zealand (41°18'S, 174°47'W) to the Northern Kermadec Trench (21°01'S, 175°12'W), involving four oceanographic stations. Both the ATACAMEX cruise with the RV *Cabo de hornos* (from January 28th to February 4st, 2018) and SO261 cruise with RV *Sonne* (March 2nd to April 2nd, 2018) were carried out in the deepest region of the trench (20°19'S — 24°15'S off northern Chile, Fig. 1, Table S1). Each sample was filtered across a hydrophilic 20-µm nylon membrane (Merck Millipore, Massachusetts, USA) and filtered onto a Sterivex fileter using a peristaltic pump. The membrane filter containing captured eDNA and cellular material from the water column was filled with lysis buffer and stored at -80°C.

### 2. Amplicon generation and sequencing

#### DNA extraction

DNA extractions were carried out by Genoscope (Évry, France) using the same protocol as described by Alberti et al. (2017)[1] for Tara Oceans water samples. The protocol is based on cryogenic grinding of membrane filters, followed by nucleic acid extraction with NucleoSpin RNA kits combined with the NucleoSpin DNA buffer set (Macherey–Nagel, Düren, Germany). A negative extraction control was performed alongside sample extractions, adding nothing in the place of sample in the first extraction step.

**PCR amplification**

Primers for PCR amplification used in this study, targeting metazoans with the COI and 18S-V1V2 loci markers.

| Locus | Target | Primers | Sequence (5'-3') | Amplicon size (bp) |
|---|---|---|---|---|
| **COI**[2] | Eukaryotes (pref. Metazoans) | mlCOIintF | GGWACWGGWTGAACWGTWTAYCCYCC | ~313 |
| | | jgHCO2198 | TAIACYTCIGGRTGICCRAARAAYCA | |
| **18S-V1V2**[3] | Eukaryotes (pref. Metazoans) | SSUF04 | GCTTGTCTCAAAGATTAAGCC | 330-390 |
| | | SSURmod | CCTGCTGCCTTCCTTRGA | |

The COI metabarcodes were generated using the mlCOIintF and jgHCO2198 primers (Table S1, Leray et al. 2013[2]). The PCR reactions (20 µL final volume) contained 2.5 ng or less of total DNA template with 0.5 µM final concentration of each primer, 3% of DMSO, 0.175 mM final concentration of dNTPs, and 1X Advantage 2 Polymerase Mix (Takara Bio, Kusatsu, Japan). Nested PCR amplifications were carried out in triplicates and consisted of an initial denaturation at 95 °C for 10 min, and 16 cycles of 10 s at 95°C, 30 s at 62 °C (−1°C per cycle), 60 s at 68 °C followed by 15 cycles of 95 °C for 10 s, 30 s at 46°C, 68 °C for 60 s, and a final extension of 68 °C for 7 min.

The 18S-V1V2 metabarcodes were generated using the SSUF04 and SSUR22mod primers (Table S1, Sinniger et al., 2016[3]) and the Phusion High Fidelity PCR Master Mix with GC buffer (ThermoFisher Scientific, Waltham, MA, USA). The PCR reactions (25 µL final volume) contained 2.5 ng or less of DNA template with 0.4 µM concentration of each primer, 3% of DMSO, and 1X Phusion Master Mix.

PCR amplifications (98 °C for 30 s; 25 cycles of 10 s at 98 °C, 30 s at 45 °C, 30 s at 72 °C; and 72 °C for 10 min) of all samples were carried out in triplicate in order to smooth the intra-sample variance while obtaining sufficient amounts of amplicons for Illumina sequencing. Amplicon triplicates were pooled and PCR products were purified using 1X AMPure XP beads (Beckman Coulter, Brea, CA, USA) cleanup. Aliquots of purified amplicons were run on an Agilent Bioanalyzer using the DNA High Sensitivity LabChip kit (Agilent Technologies, Santa Clara, CA, USA) to check their lengths, and quantified with a Qubit fluorometer (Invitrogen, Carlsbad, CA, USA).

**Clean-up**

The PCR products were pooled and cleaned using 1X AMPure XP beads, and amplicon lengths were checked with the DNA High Sensitivity LabChip kit (Agilent Technologies, Santa Clara, CA, USA). Subsequently the concentration of the purified PCR products was quantified with a Qubit fluorometer (Invitrogen, Carlsbad, CA, USA).

**Amplicon library preparation**

From each purified PCR product pool, one hundred ng were end-repaired, A-tailed and ligated to Illumina adapters on a Biomek FX Laboratory Automation Workstation (Beckman Coulter, Brea, CA, USA). Afterwards, each library was amplified using a Kapa Hifi HotStart NGS library Amplification kit (Kapa Biosystems, Wilmington, MA, USA) and purified again with 1X AMPure XP beads.

**Sequencing library quality control**

Libraries were quantified with both a Quant-iT dsDNA HS assay kits using a Fluoroskan Ascent microplate fluorometer (Thermo Fisher Scientific, Waltham, MA, USA) and qPCR with the KAPA Library Quantification Kit for Illumina Libraries (Kapa Biosystems, Wilmington, MA, USA) on a MxPro instrument (Agilent Technologies, Santa Clara, CA, USA). A high-throughput microfluidic capillary electrophoresis system (LabChip GX, Perkin Elmer, Waltham, MA, USA) was used to assess the library profiles.

**Sequencing procedures**

The concentrations of all libraries were normalized to 10 nM by addition of 10 mM Tris-Cl (pH 8.5) and clusters generated according to the Illumina Cbot User Guide (Part # 15006165). Paired-end sequencing (2×250bp) of the libraries was performed on either the HiSeq 4000 or HiSeq 2500 instruments (Illumina, San Diego, CA, USA). In order to enhance the sequence quality and counteracting the low contrast of the first few bp due to adapters and primers, the loading concentration of the libraries was reduced from 12-14 pM to 8-9 pM, while PhiX DNA spike-in was increased (20% instead of 1%). Sequencing was otherwise performed according to the HiSeq 4000 System User Guideline (Part # 15011190) and the HiSeq 2500 System User Guideline (Part # 15035786).

### 3. Sequence analysis

**COI sequence datasets**: The following is a sketch of the sequence analysis of the COI metabarcode sequences:

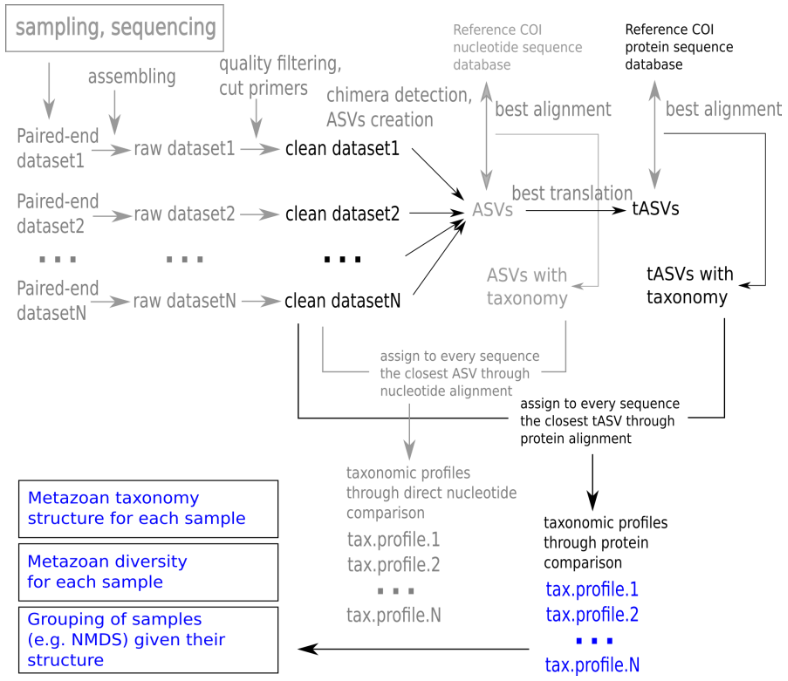


Paired-end reads were assembled and quality filtered with vsearch[4] (version 2.17.1), using the parameters maxEE of 0.5, minimum length of 250 bp, and maximum length of 500 bp. The primers in the resulting metabarcoding sequences were detected removed with pTrimmer[5] (version 1.3.4). The sequences lacking the primers were discarded. All the sequences from all the datasets were then merged into a single file and the unique sequences were discriminated with vsearch. Chimeras were then removed with the same program and in the same process, the Amplicon Sequence Variants (ASVs) were created. To reduce complexity and to establish yet another filtering procedure these ASVs were then translated into protein sequences (translated ASV or tASV), using the longest translation of the six open reading frames using the 33 translation tables (genetic codes) known to date. Since the COI region amplified by the primers has an approximate size of 300 bp, the tASV shorter than 90 amino acids were discarded. The retained tASVs were aligned against a custom reference sequence database of COI sequences with BLASTP (bitscore cutoff of 50). This reference COI sequences database was constructed with the COI protein sequences from the NCBI mitochondrial RefSeq database[6] plus the complete COI sequences from the COI-arbitrator database[7] and the bacterial COX1/COI sequences from the COG0843 in the COG database[8]. The bacterial sequences were included because it has been reported that the metazoan COI primers could potentially amplify bacterial sequences[9], and with this procedure they were identified and discarded for further analysis of metazoa. The sequences from all the datasets were then compared (through BLASTX, with a bitscore cutoff of 50) with the tASVs database and assigned taxonomy accordingly to construct the taxonomic profiles for each sample.

**18S V1V2 sequence datasets**: The procedure for these datasets was similar to the previously described for the COI sequence datasets, except that in this case the ASVs were directly compared as nucleotide sequences using BLASTN (parameter task dc-megablast, bitscore cutoff of 50) against the SILVA database[10] (version 138.1), to construct the corresponding taxonomic profiles.

**Phylogenetic analyses**: The phylogenetic trees of 18S ASVs in the Figures S5, S6 and S7 were constructed as follows. First, reference sequences for hydrozoans, copepods and tunicates were collected from the literature and from respective searches in Genbank, generating separate datasets for each taxonomic group (Tables S5, S6 and S7). These datasets were complemented with the ASVs sequences associated with each taxonomic group according to the previously described procedure (comparison with the SILVA database). Second, these datasets of sequences were then aligned with mafft[11] with parameters for high accuracy (defined in the documentation of this software). The respective alignments were used to create phylogenetic trees with IQ-TREE2[12] with paratemeters (-nt AUTO, -ntop 50, -nbest 10, -nstop 1000 and -bb 1000). Third, the resulting consensus phylogenetic trees (Newick format) were edited with Bosque[13] (available at https://inf.imo-chile.cl/software/bosque.html) and Inkscape.

### 4. Supplementary figures

**Figure S1**. Taxonomic composition in terms of presence/absence and relative abundances of the 18S ASVs and COI translated ASVs. The names of the samples are split into three parts, encoding the name of the cruise/station, the water filter size range ("02" for the range 0.2-3 µm, and "3" for the range 3-20 µm), and the depth (meters), respectively. More details about the samples in Table S1. Note that ctenophores and tunicates were not detected with the COI metabarcodes, while placozoans and Porifera were detected in COI but not in the 18S data. These differences most likely reflect the differential taxonomic affinity of the primers already reported in the literature[14,15].

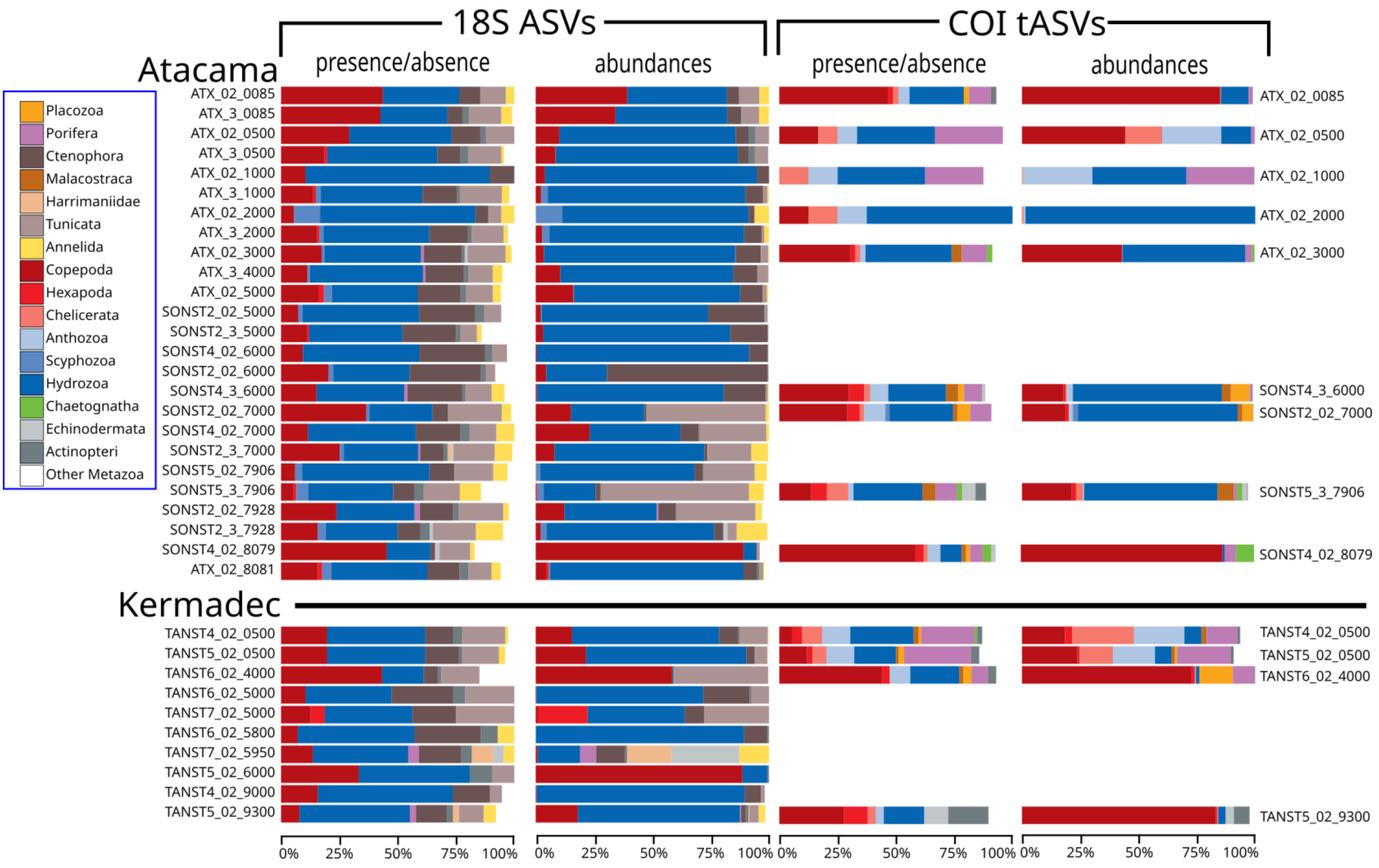

**Figure S2**. Taxonomic distribution of the 18S ASVs (presence/absence) associated with hydrozoans (a) and copepods (b) at the family taxonomic rank. The names of the samples are split into three parts, encoding the name of the cruise/station, the water filter size range ("02" for the range 0.2-3 μm, and "3" for the range 3-20 μm), and the depth (meters). More details about the samples in Table S1.

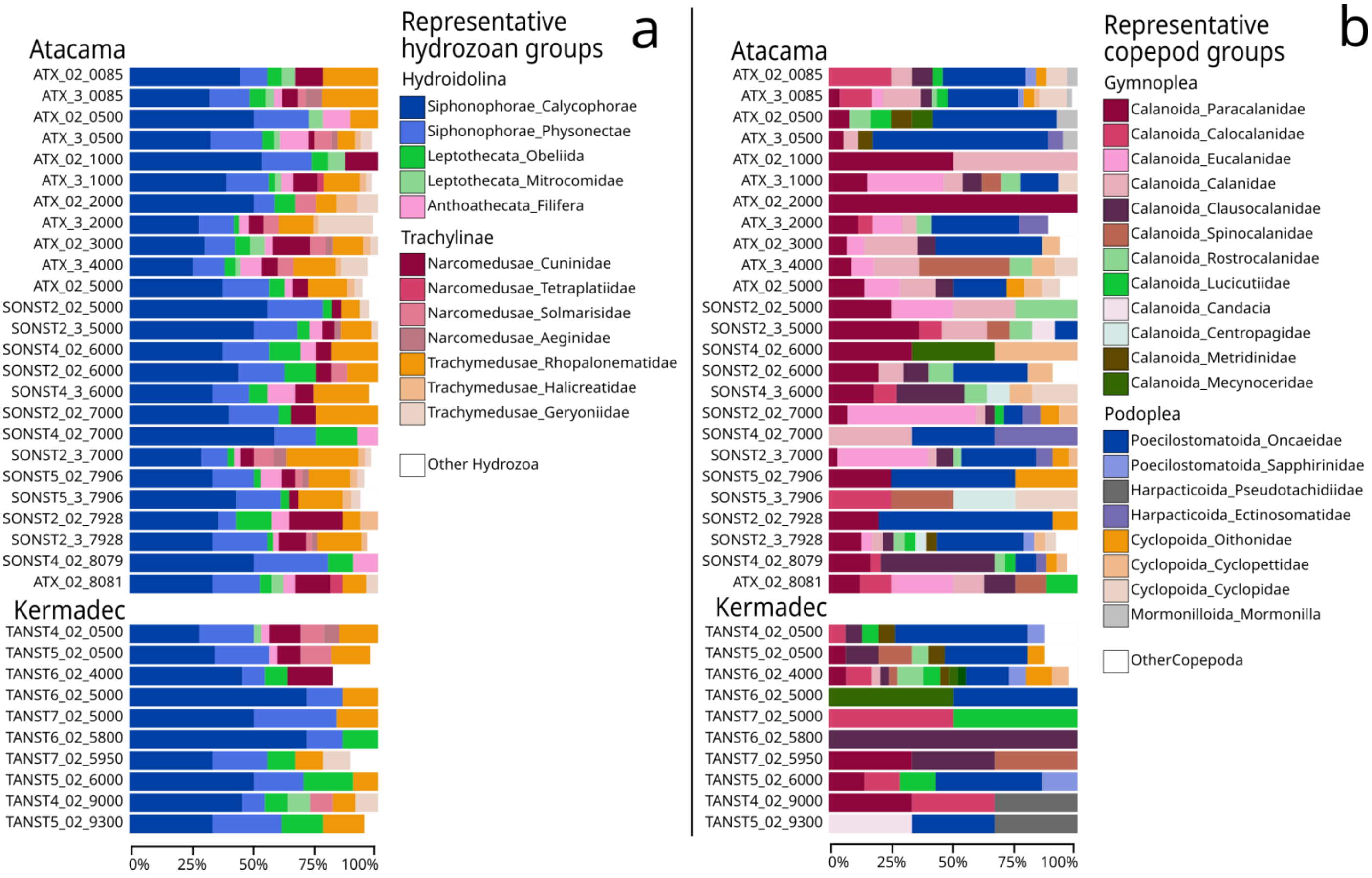

**Figure S3**. Taxonomic distribution of the 18S ASVs (relative abundances) associated with hydrozoans (a) and copepods (b) at the family taxonomic rank. The names of the samples are split into three parts, encoding the name of the cruise/station, the water filter size range ("02" for the range 0.2-3 µm, and "3" for the range 3-20 µm), and the depth (meters). More details about the samples in Table S1.

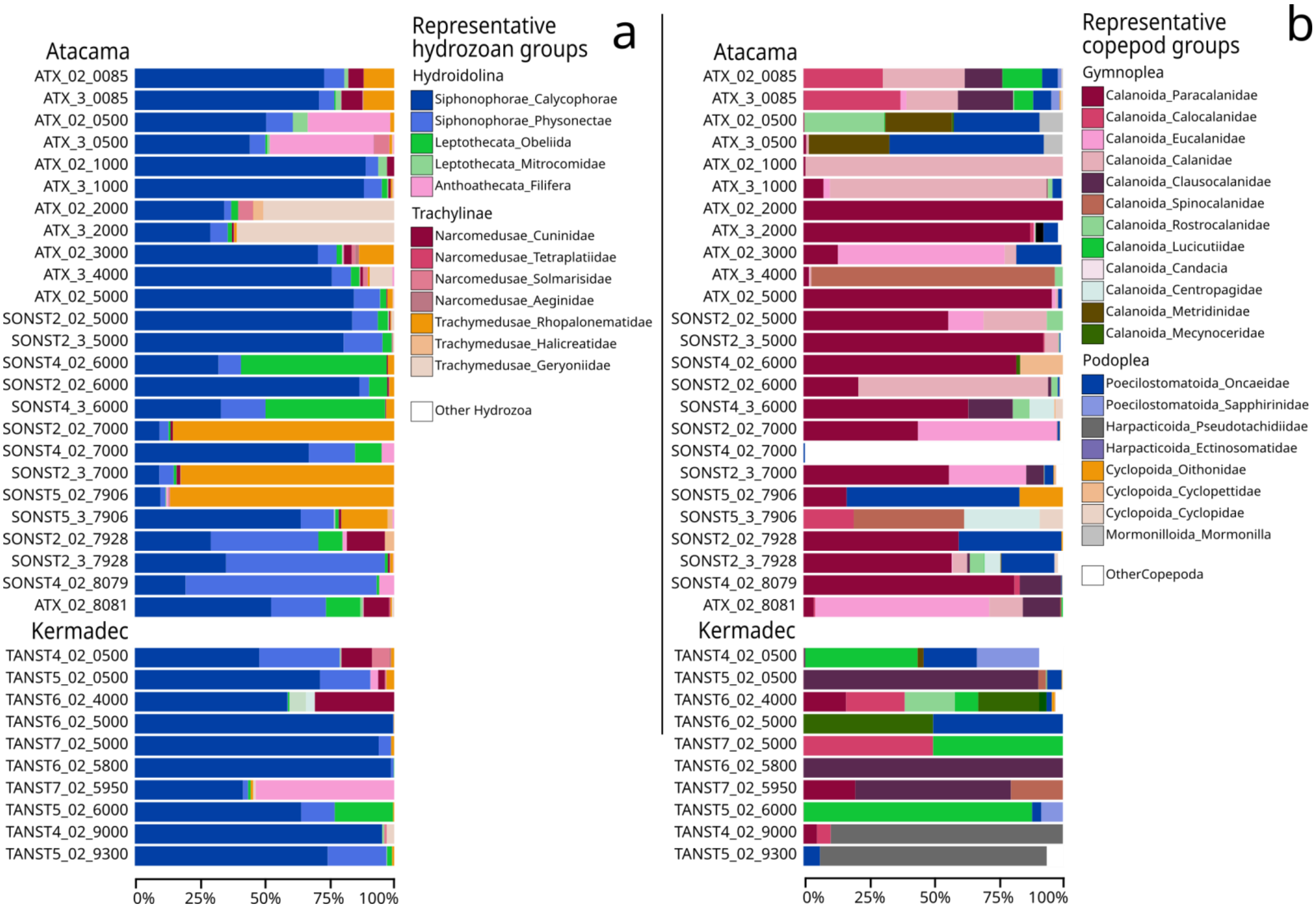

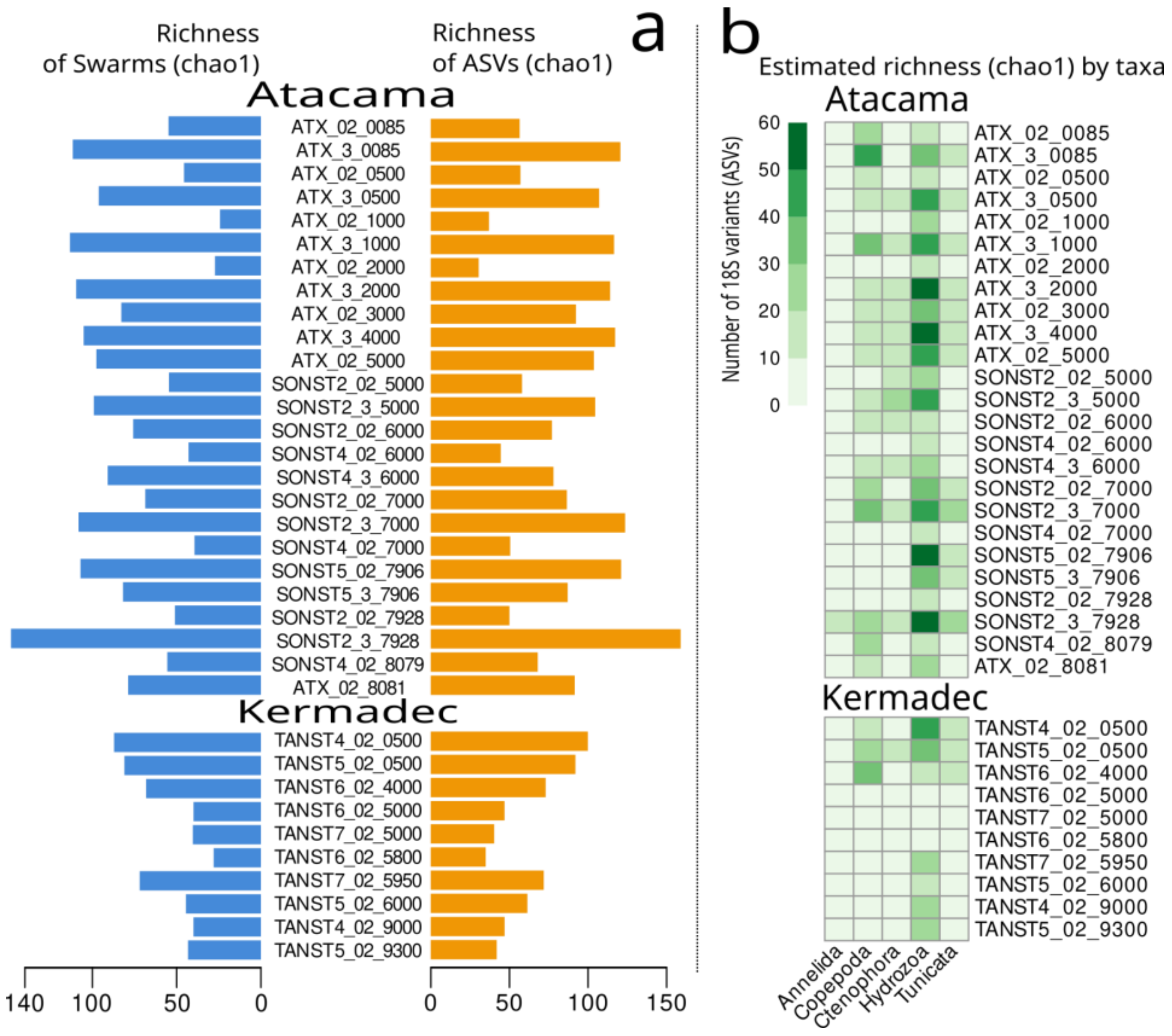


**Figure S4**. **a)** Metazoan genetic richness per sample (estimated by chao1) of the “swarm” clusters (left) and ASVs (right), obtained from the 18S metabarcoding sequences. The pattern of genetic richness in relation to depth is very similar between “swarm” clusters and ASVs and only differs in amplitude. **b)** Metazoan genetic richness (estimated by chao1) of the 18S metabarcoding sequences in the most representative taxonomic groups for each sample.

**Figure S5**. **a)** Metazoan richness and diversity of the translated metabarcode COI sequences per sample. The diversity was estimated by the Gini-Simpson index (left) and the richness was estimated by the Chao1 index (red bar under the observed richness, right). **b)** Metazoan richness (Chao1) of the translated metabarcode COI sequences in the most relatively abundant taxonomic groups (Fig. 1) for each sample. The nomenclature of the sample names is described in Figure 1.

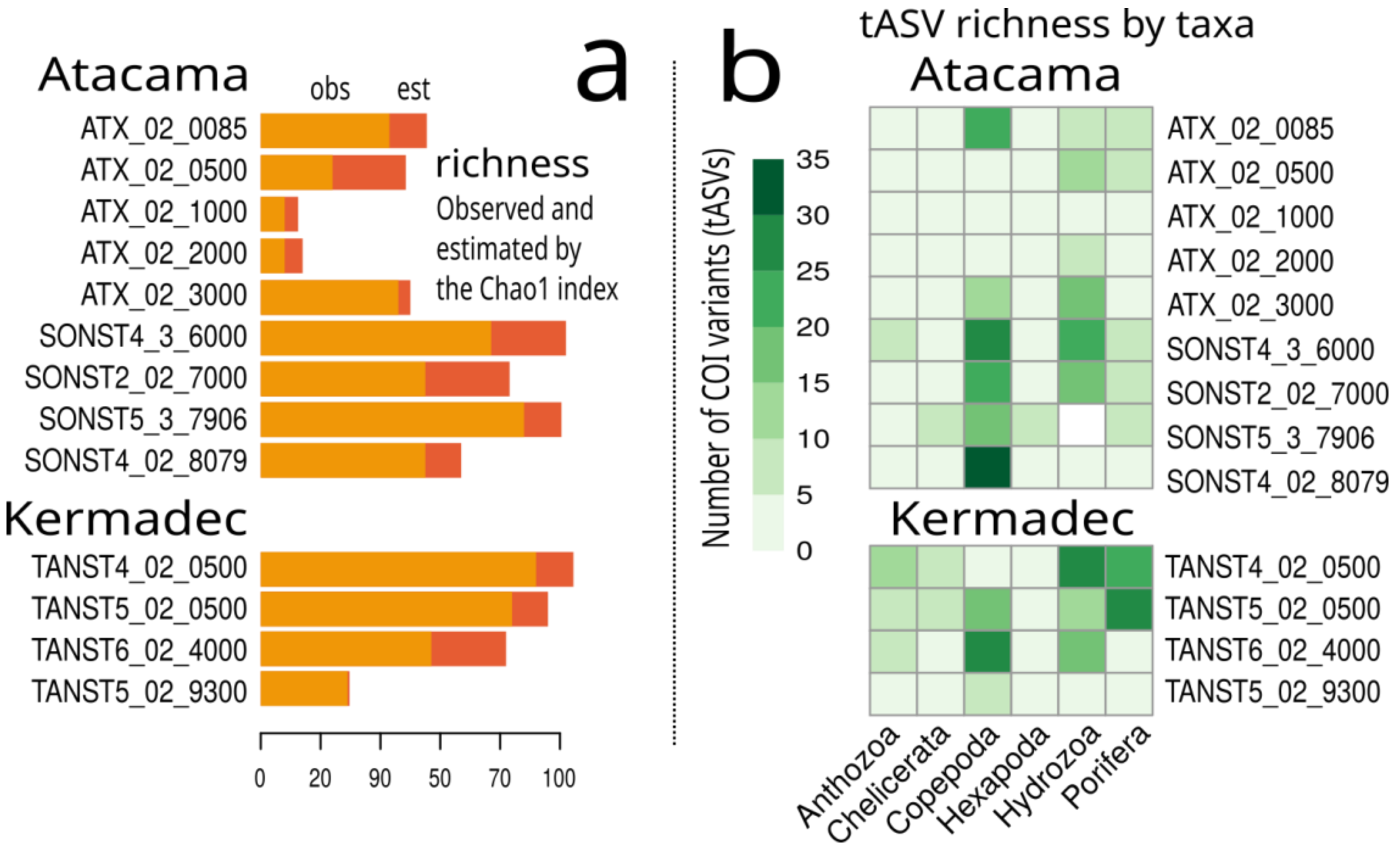

**Figure S6**. Phylogenetic subtrees of 18S ASVs associated with hydrozoan orders with clades exclusively composed by ASVs generated in this study (general phylogenetic tree shown in Fig. 4a). The ASV sequences are labeled according to their occurrences in the samples from Atacama (A) and Kermadec (K). Additionally, this label includes a code to indicate whether the ASV occurred only in the samples <4000 m depth (‘n’), in those > 4000 m depth (‘d’, abbyso-hadal), or if it was found in both partitions (‘u’). Thus, an ASV found only > 4000m in Atacama is labeled ‘Ad’, whereas an ASV found in both systems only > 4000 m is labeled ‘AdKd’, and so on. The statistical support numbers in the branches are UFBoot values and thus they can directly be interpreted as approximate probabilities. The references sequences used in this tree can be found in Table S5. The subtrees correspond to hydroazoan orders: **a)** Siphonophores, **b)** Rhopalonematidae, **c)** Geryonidae and, **d)** Clytiidae.

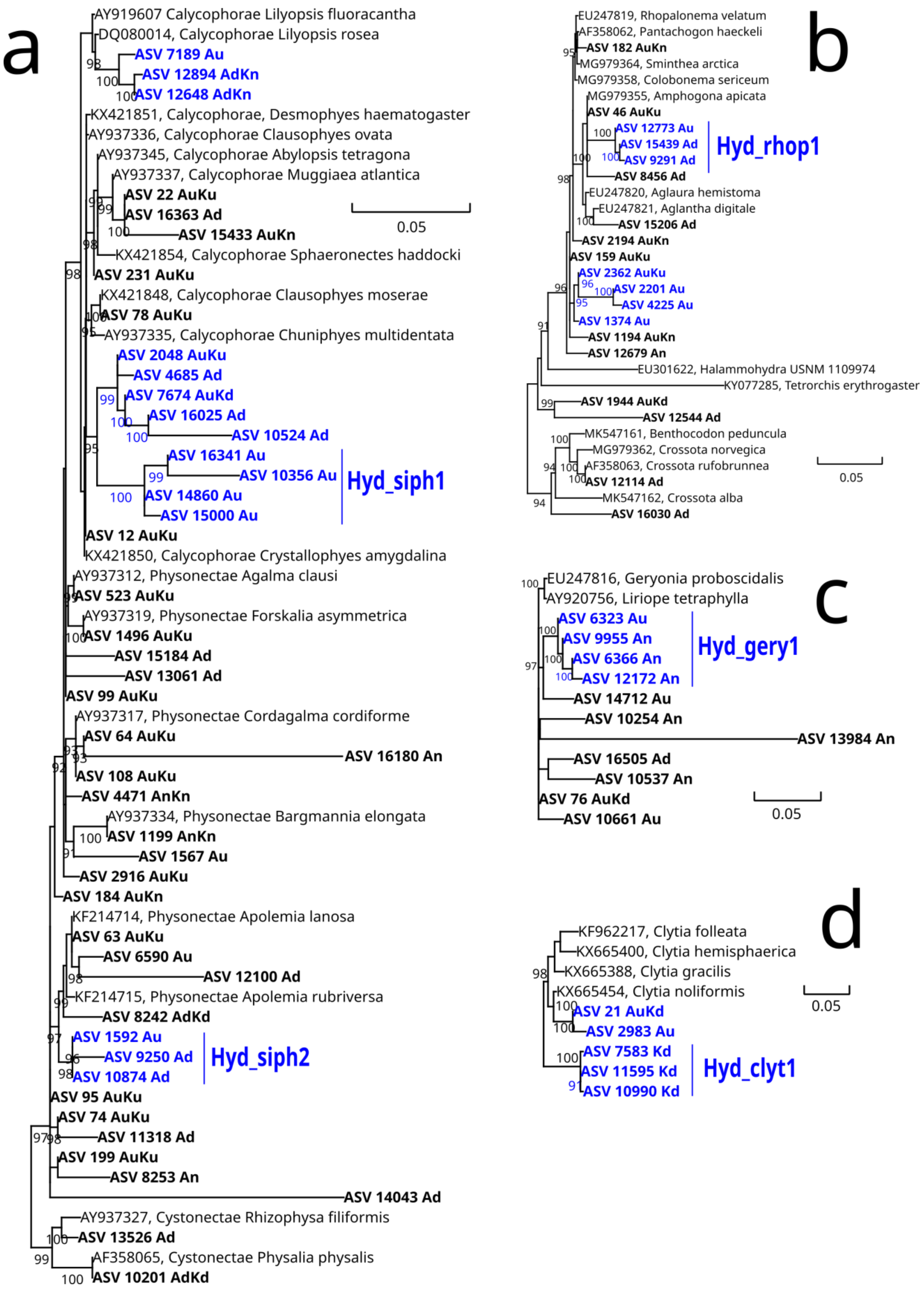

**Figure S7**. Phylogenetic subtrees of 18S ASVs associated with copepod orders with clades exclusively composed by ASVs generated in this study (general phylogenetic tree shown in Fig. 4b). The ASV sequences are labeled according to their occurrences in the samples from Atacama (A) and Kermadec (K). Additionally, this label includes a code to indicate whether the ASV occurred only in the samples <4000 m depth ('n'), in those > 4000 m depth ('d', abbyso-hadal), or if it was found in both partitions ('u'). The statistical support numbers in the branches are UFBoot values and thus they can directly be interpreted as approximate probabilities. The references sequences used in this tree can be found in Table S6. The subtrees correspond to copepod families: **a)** Oncaeidae, **b)** Clausocalanidae, and **c)** Calanidae (genus cosmocalanus).

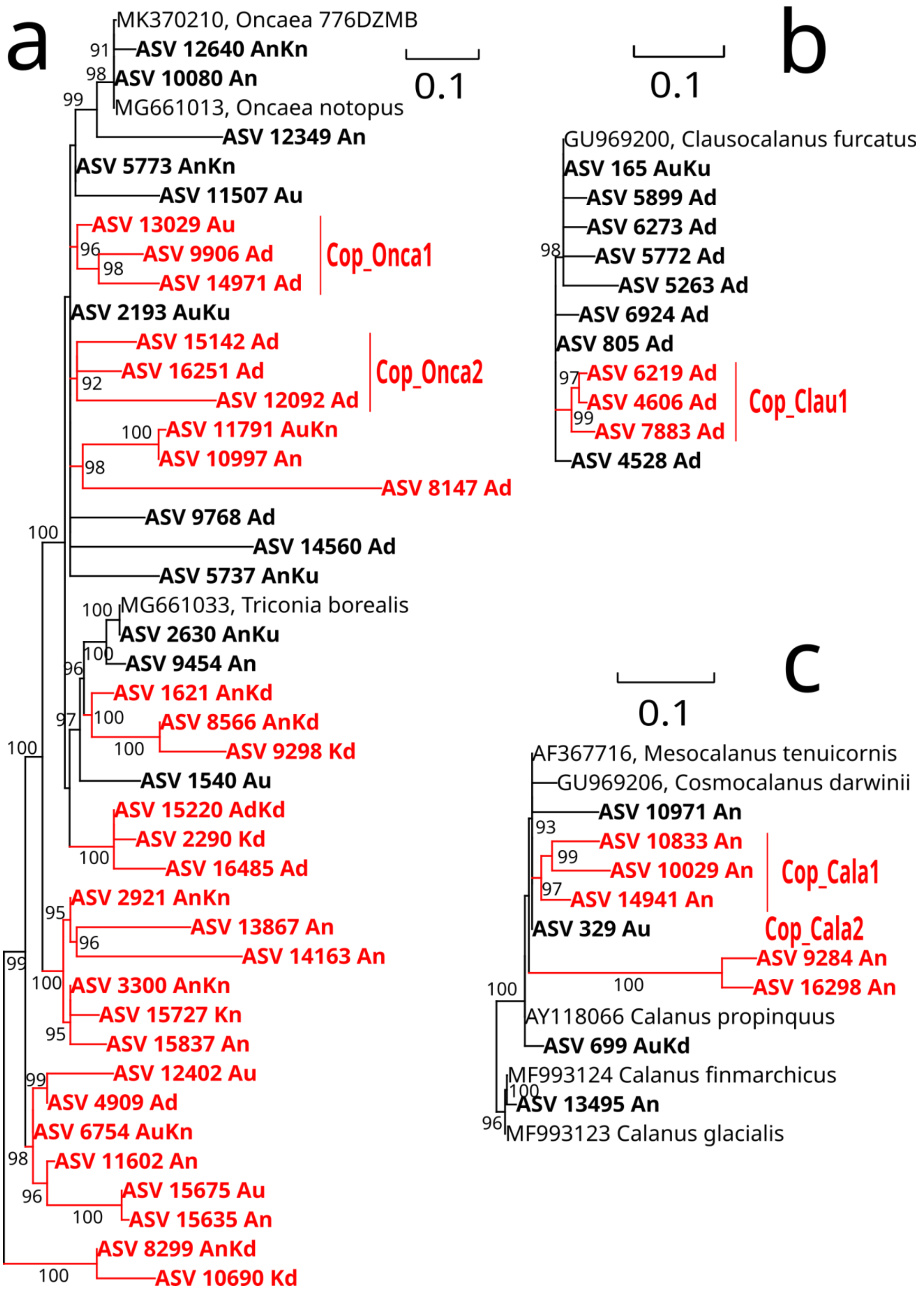

**Figure S8**. Phylogenetic tree of the 18S ASVs associated with tunicates. The ASV sequences are labeled according to their occurrences in the samples from Atacama (A) and Kermadec (K). Additionally, this label includes a code to indicate whether the ASV occurred only in the samples <4000 m depth (‘n’), in those > 4000 m depth (‘d’, abbyso-hadal), or if it was found in both partitions (‘u’). The whole set of reference sequences utilized for these trees can be found in Table S7.

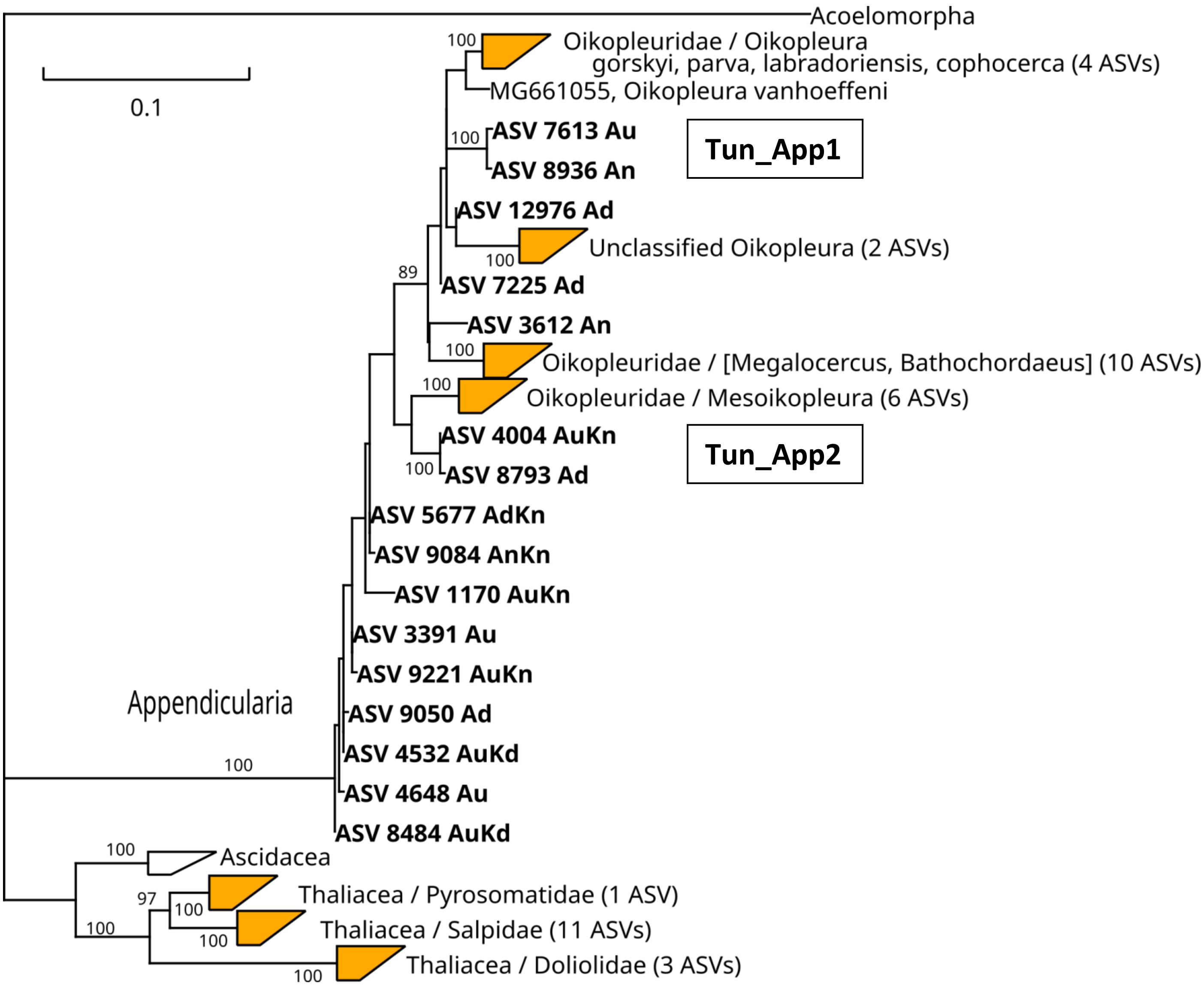

### Supplementary tables

**Table S1**. Sampling locations, filtering conditions and target genes for the metabarcoding analysis. The column named as “Filter” refers to the free living fraction (FL, between 0.2 and 3um) and particulate fraction (PA, between 3 and 20um).

| Sample name | Cruise | Trench | St. | Lat. | Lon. | Date | Time | Depth (m) | Filter | Rosette type | Water (L) | Target genes |
|---|---|---|---|---|---|---|---|---|---|---|---|---|
| ATX_02_0085 | ATACAMEX | Atacama | 1 | -23°14' | -71°11' | January 30, 2018 | 20:18 | 85 | FL | CTD | 40 | 18S,COI |
| ATX_3_0085 | ATACAMEX | Atacama | 1 | -23°14' | -71°11' | January 30, 2018 | 20:18 | 85 | PA | CTD | 40 | 18S |
| ATX_02_0500 | ATACAMEX | Atacama | 1 | -23°14' | -71°11' | January 30, 2018 | 20:18 | 500 | FL | CTD | 40 | 18S,COI |
| ATX_3_0500 | ATACAMEX | Atacama | 1 | -23°14' | -71°11' | January 30, 2018 | 21:18 | 500 | PA | CTD | 40 | 18S |
| ATX_02_1000 | ATACAMEX | Atacama | 1 | -23°14' | -71°11' | January 30, 2018 | 22:18 | 1000 | FL | CTD | 40 | 18S,COI |
| ATX_3_1000 | ATACAMEX | Atacama | 1 | -23°14' | -71°11' | January 30, 2018 | 23:18 | 1000 | PA | CTD | 40 | 18S |
| ATX_02_2000 | ATACAMEX | Atacama | 1 | -23°14' | -71°11' | January 30, 2018 | 0:18 | 2000 | FL | CTD | 40 | 18S,COI |
| ATX_3_2000 | ATACAMEX | Atacama | 1 | -23°14' | -71°11' | January 30, 2018 | 1:18 | 2000 | PA | CTD | 40 | 18S |
| ATX_02_3000 | ATACAMEX | Atacama | 1 | -23°14' | -71°11' | January 30, 2018 | 2:18 | 3000 | FL | CTD | 40 | 18S,COI |
| ATX_3_4000 | ATACAMEX | Atacama | 1 | -23°14' | -71°11' | January 30, 2018 | 3:18 | 4000 | PA | CTD | 40 | 18S |
| ATX_02_5000 | ATACAMEX | Atacama | 1 | -23°14' | -71°11' | January 30, 2018 | 4:18 | 5000 | FL | CTD | 40 | 18S |
| SONST2_02_5000 | SONNE | Atacama | 2 | -21°28' | -71°7' | March 22, 2018 | 21:03 | 5000 | FL | CTD | 15.19 | 18S |
| SONST2_3_5000 | SONNE | Atacama | 2 | -21°28' | -71°7' | March 22, 2018 | 22:03 | 5000 | PA | CTD | 15.19 | 18S |
| SONST2_02_6000 | SONNE | Atacama | 2 | -21°28' | -71°7' | March 22, 2018 | 23:03 | 6000 | FL | CTD | 23.4 | 18S |
| SONST4_02_6000 | SONNE | Atacama | 4 | -23°12' | -71°12' | March 13, 2018 | 10:36 | 6000 | FL | CTD | 42.6 | 18S |
| SONST4_3_6000 | SONNE | Atacama | 4 | -23°12' | -71°12' | March 13, 2018 | 11:36 | 6000 | PA | CTD | 42.6 | 18S,COI |
| SONST2_02_7000 | SONNE | Atacama | 2 | -21°28' | -71°7' | March 24, 2018 | 23:05 | 7000 | FL | HR | 25.1 | 18S,COI |
| SONST2_3_7000 | SONNE | Atacama | 2 | -21°28' | -71°7' | March 24, 2018 | 23:05 | 7000 | PA | HR | 25.1 | 18S |

| | | | | | | | | | | | | |
|---|---|---|---|---|---|---|---|---|---|---|---|---|
| SONST4_02_7000 | SONNE | Atacama | 4 | -23°12' | -71°12' | March 13, 2018 | 23:22 | 7000 | FL | HR | 25.3 | 18S |
| SONST5_02_7906 | SONNE | Atacama | 5 | -23°29' | -71°13' | March 11, 2018 | 22:32 | 7906 | FL | NL | 44.9 | 18S,COI |
| SONST5_3_7906 | SONNE | Atacama | 5 | -23°29' | -71°13' | March 11, 2018 | 22:32 | 7906 | PA | NL | 44.9 | 18S,COI |
| SONST2_02_7928 | SONNE | Atacama | 2 | -21°28' | -71°7' | March 22, 2018 | 12:42 | 7928 | FL | NL | 20.6 | 18S |
| SONST2_3_7928 | SONNE | Atacama | 2 | -21°28' | -71°7' | March 22, 2018 | 12:42 | 7928 | PA | NL | 20.6 | 18S |
| SONST4_02_8079 | SONNE | Atacama | 4 | -23°12' | -71°12' | March 14, 2018 | 18:00 | 8079 | FL | NL | 12.9 | 18S,COI |
| ATX_02_8081 | ATACAMEX | Atacama | 1 | -23°14' | -71°11' | February 1, 2018 | 9:55 | 8081 | FL | NL | 50 | 18S |
| TANST4_02_0500 | HADES-ERC | Kermadec | 4 | -31°08' | -176°48' | December 8, 2017 | 14:50 | 500 | FL | CTD | 31 | 18S,COI |
| TANST5_02_0500 | HADES-ERC | Kermadec | 5 | -31°55' | -177°17' | December 10, 2017 | 1:15 | 500 | FL | CTD | 18 | 18S,COI |
| TANST6_02_4000 | HADES-ERC | Kermadec | 6 | -32° 09' | 177°23' | December 1, 2017 | 2:45 | 4000 | FL | CTD | 18 | 18S,COI |
| TANST6_02_5000 | HADES-ERC | Kermadec | 6 | -32° 09' | 177°23' | December 1, 2017 | 2:45 | 5000 | FL | CTD | 18.6 | 18S |
| TANST7_02_5000 | HADES-ERC | Kermadec | 7 | -32°10' | 176°32' | November 28, 2017 | 21:30 | 5000 | FL | CTD | 16 | 18S |
| TANST6_02_5800 | HADES-ERC | Kermadec | 6 | -32° 09' | 177°23' | December 1, 2017 | 2:45 | 5800 | FL | CTD | 18.7 | 18S |
| TANST7_02_5950 | HADES-ERC | Kermadec | 7 | -32°10' | 176°32' | November 28, 2017 | 21:30 | 5950 | FL | CTD | 10 | 18S |
| TANST5_02_6000 | HADES-ERC | Kermadec | 5 | -31°55' | -177°17' | December 10, 2017 | 1:15 | 6000 | FL | CTD | 18 | 18S |
| TANST4_02_9000 | HADES-ERC | Kermadec | 4 | -31°08' | -176°48' | December 8, 2017 | 0:25 | 9000 | FL | HR | 28 | COI |
| TANST5_02_9300 | HADES-ERC | Kermadec | 5 | -31°55' | -177°17' | December 10, 2017 | 9:40 | 9300 | FL | HR | 17.2 | 18S,COI |

**Table S2.** Number of COI tASVs per metazoan phyla obtained in Atacama. The second and third columns present the biome ranges in which the respective phylum has been documented in the current literature. The fourth column corresponds to the number ASVs exclusively found in the samples above 4000m depth, whereas the fifth column shows the number of ASVs exclusively found in the samples below 4000m depth. The sixth column shows the ASVs that were found both above and below 4000m.

| Phylum | Aquatic Biomes | General Biomes | <4000m | >4000m | Ubiq. | Total |
|---|---|---|---|---|---|---|
| **Annelida (mostly polychaetes)** | Pelagic/Benthic | Marine/Freshwater/Terrestrial | 0 | 0 | 0 | 0 |
| **Arthropoda (mostly copepods)** | Pelagic/Benthic | Marine/Freshwater/Terrestrial | 14 | 53 | 25 | 92 |
| **Brachiopoda** | Benthic | Marine | 0 | 0 | 0 | 0 |
| **Bryozoa** | Pelagic/Benthic | Marine/Freshwater | 0 | 0 | 0 | 0 |
| **Chaetognatha** | Pelagic/Benthic | Marine | 0 | 1 | 1 | 2 |
| **Chordata (mostly tunicates)** | Pelagic/Benthic | Marine/Freshwater/Terrestrial | 1 | 8 | 0 | 9 |
| **Cnidaria (mostly hydrozoans)** | Pelagic/Benthic | Marine/Freshwater | 20 | 37 | 18 | 75 |
| **Ctenophora** | Pelagic/Benthic | Marine/Freshwater | 9 | 7 | 6 | 22 |
| **Cycliophora** | Benthic | Marine | 0 | 0 | 0 | 0 |
| **Echinodermata** | Benthic | Marine | 0 | 6 | 0 | 6 |
| **Entoprocta** | Pelagic/Benthic | Marine/Freshwater | 0 | 0 | 0 | 0 |
| **Gastrotricha** | Benthic | Marine/Freshwater | 0 | 0 | 0 | 0 |
| **Gnathostomulida** | Benthic | Marine | 0 | 0 | 0 | 0 |
| **Hemichordata** | Pelagic/Benthic | Marine | 0 | 0 | 0 | 0 |
| **Micrognathozoa** | Pelagic/Benthic | Freshwater | 0 | 0 | 0 | 0 |
| **Mollusca** | Pelagic/Benthic | Marine/Freshwater/Terrestrial | 0 | 3 | 1 | 4 |
| **Nematoda** | Benthic | Marine/Freshwater/Terrestrial | 0 | 0 | 0 | 0 |
| **Nemertea** | Benthic | Marine/Freshwater/Terrestrial | 0 | 0 | 0 | 0 |
| **Onychophora** | NA | Terrestrial | 0 | 0 | 0 | 0 |
| **Phronida** | Benthic | Marine | 0 | 0 | 0 | 0 |
| **Placozoa** | Pelagic/Benthic | Marine | 0 | 2 | 1 | 3 |
| **Platyhelminthes** | Pelagic/Benthic | Marine | 0 | 0 | 0 | 0 |
| **Porifera** | Pelagic/Benthic | Marine/Freshwater | 9 | 7 | 6 | 22 |
| **Priapulida** | Benthic | Marine | 0 | 0 | 0 | 0 |
| **Syndermata** | Pelagic/Benthic | Marine/Freshwater | 0 | 0 | 0 | 0 |
| **Tardigrada** | Pelagic/Benthic | Marine/Freshwater/Terrestrial | 0 | 0 | 0 | 0 |
| **Xenacoelomorpha** | Pelagic/Benthic | Marine | 0 | 0 | 0 | 0 |

**Table S3.** Taxonomic affiliation at the family level (when possible) of the 18S ASVs from iconic marine phyla other than Copepoda, Cnidaria and the subphylum Tunicate within Chordata in Atacama.

| ***Simplified taxonomic lineage*** | ***ASVs*** |
|---|---|
| Annelida;Polychaeta;Echiura;Echiuroidea;Echiuridae;Thalassematinae | 1 |
| Annelida;Polychaeta;Palpata;Aciculata;Phyllodocida;Glyceridae | 1 |
| Annelida;Polychaeta;Palpata;Aciculata;Phyllodocida;Phyllodocidae | 20 |
| Annelida;Polychaeta;Palpata;Aciculata;Phyllodocida;Typhloscolecidae | 2 |
| Annelida;Polychaeta;Palpata;Canalipalpata;Flabelligerida;Flabelligeridae | 1 |
| Annelida;Polychaeta;Palpata;Canalipalpata;Sabellida;Serpulidae | 1 |
| Annelida;Polychaeta;Scolecida;Opheliidae | 1 |
| Annelida;Polychaeta;Scolecida;Spionida;Spionidae | 1 |
| Annelida;Polychaeta;Sipuncula;...Phascolosomatidae | 1 |
| Arthropoda;Chelicerata;Arachnida;Acari;Acariformes | 1 |
| Arthropoda;Crustacea;Branchiopoda;...Cyclestheriidae | 1 |
| Arthropoda;Crustacea;Malacostraca;...Euphausiidae | 2 |
| Arthropoda;Crustacea;Malacostraca;...Mysidae | 1 |
| Arthropoda;Crustacea;Malacostraca;Decapoda;...Pinnotheridae | 1 |
| Arthropoda;Crustacea;Malacostraca;Decapoda;...Xanthidae | 1 |
| Arthropoda;Crustacea;Oligostraca;Ostracoda;...Halocyprididae | 6 |
| Arthropoda;Hexapoda;Insecta;Dicondylia;Pterygota;Neoptera | 6 |
| Chaetognatha;Sagittoidea;...Eukrohniidae | 7 |
| Chaetognatha;Sagittoidea;...Sagittidae | 10 |
| Chordata;...Actinopteri;...Euteleosteomorpha;...Acanthomorphata;...Atherinidae | 1 |
| Chordata;...Actinopteri;...Euteleosteomorpha;...Acanthomorphata;...Carangidae | 1 |
| Chordata;...Actinopteri;...Euteleosteomorpha;...Acanthomorphata;...Cyprinodontidae | 1 |
| Chordata;...Actinopteri;...Euteleosteomorpha;...Acanthomorphata;...Gobiidae | 1 |
| Chordata;...Actinopteri;...Euteleosteomorpha;...Acanthomorphata;...Percichthyidae | 2 |
| Chordata;...Actinopteri;...Euteleosteomorpha;...Acanthomorphata;...Sebastidae | 2 |
| Chordata;...Actinopteri;...Euteleosteomorpha;...Retropinnidae | 4 |
| Chordata;...Actinopteri;...Otomorpha;...Clupeidae | 2 |
| Chordata;...Actinopteri;...Otomorpha;...Cyprinidae | 1 |
| Ctenophora;Tentaculata;Cydippida;Dryodoridae | 2 |
| Ctenophora;Tentaculata;Cydippida;Haeckeliidae | 4 |
| Ctenophora;Tentaculata;Cydippida;Mertensiidae | 13 |
| Ctenophora;Tentaculata;Cydippida;Pleurobrachiidae | 3 |
| Ctenophora;Tentaculata;Lobata;Lobatolampeidae | 1 |
| Ctenophora;Tentaculata;Lobata;Ocyropsidae | 2 |
| Ctenophora;Tentaculata;Lobata;unclassified_Lobata | 5 |
| Ctenophora;Tentaculata;Platyctenida;Coeloplanidae | 3 |
| Echinodermata;Echinozoa;...Elpidiidae | 3 |
| Echinodermata;Echinozoa;...Myriotrochidae | 2 |
| Hemichordata;Enteropneusta;...Tergivelum_baldwinae | 1 |
| Hemichordata;Enteropneusta;Harrimaniidae | 2 |
| Mollusca;Bivalvia;Heterodonta;...Lucinidae | 1 |
| Mollusca;Gastropoda;Caenogastropoda;...Fasciolariidae | 1 |
| Mollusca;Gastropoda;Caenogastropoda;...Muricidae | 1 |
| Mollusca;Gastropoda;Heterobranchia;...Cavoliniidae | 2 |
| Mollusca;Gastropoda;Heterobranchia;...Limacinidae | 1 |
| Porifera;Calcarea;Calcinea;Clathrinida;Leucettidae | 1 |

| | |
|---|---|
| Porifera;Calcarea;Calcinea;Murrayonida;Lelapiellidae | 1 |
| Porifera;Demospongiae;Heteroscleromorpha;Axinellida;Raspailiidae | 1 |
| Porifera;Demospongiae;Heteroscleromorpha;Poecilosclerida;Hymedesmiidae | 1 |

**Table S4.** Taxonomic affiliation at the family level (when possible) of the 18S ASVs from iconic marine phyla other than Copepoda, Cnidaria and the subphylum Tunicate within Chordata in Kermadec.

| ***Simplified taxonomic lineage*** | ***ASVs*** |
|---|---|
| Annelida;Polychaeta;Palpata;Aciculata;Phyllodocida;Phyllodocidae | 2 |
| Annelida;Polychaeta;Palpata;Aciculata;Phyllodocida;Typhloscolecidae | 1 |
| Annelida;Polychaeta;Polychaeta_incertae_sedis;Nerillidae | 1 |
| Annelida;Polychaeta;Scolecida;Capitellida;Scalibregmatidae | 1 |
| Annelida;Polychaeta;Scolecida;Spionida;Cirratulidae | 1 |
| Arthropoda;Crustacea;Hexanauplia;Thecostraca;...Peltogastridae | 1 |
| Arthropoda;Crustacea;Malacostraca;Eumalacostraca;...Euphausiidae | 1 |
| Arthropoda;Hexapoda;Insecta;Dicondylia;Pterygota;Neoptera | 1 |
| Arthropoda;Myriapoda;Diplopoda;Helminthomorpha;Sphaerotheriida;Sphaerotheriidae | 1 |
| Chaetognatha;Sagittoidea;Aphragmophora;Ctenodontina;Sagittidae | 2 |
| Chordata;...Actinopteri;...Euteleosteomorpha;...Acanthomorphata;...Atherinomorphae | 1 |
| Chordata;...Actinopteri;...Euteleosteomorpha;...Acanthomorphata;...Pomacanthidae | 1 |
| Chordata;...Actinopteri;...Euteleosteomorpha;...Acanthomorphata;...Sebastidae | 1 |
| Chordata;...Actinopteri;...Euteleosteomorpha;Stomiatii;Osmeriformes;Retropinnidae | 3 |
| Chordata;...Actinopteri;...Otomorpha;Clupei;Clupeiformes;Clupeoidei;Clupeidae | 2 |
| Ctenophora;Tentaculata;Cydippida;Dryodoridae | 2 |
| Ctenophora;Tentaculata;Cydippida;Mertensiidae | 8 |
| Ctenophora;Tentaculata;Cydippida;Pleurobrachiidae | 1 |
| Ctenophora;Tentaculata;Lobata;Lobatolampeidae | 1 |
| Ctenophora;Tentaculata;Lobata;Ocyropsidae | 1 |
| Ctenophora;Tentaculata;Lobata;unclassified_Lobata | 3 |
| Ctenophora;Tentaculata;Platyctenida;Coeloplanidae | 2 |
| Echinodermata;Echinozoa;Holothuroidea;Aspidochirotacea;Elasipodida;Elpidiidae | 1 |
| Hemichordata;Enteropneusta;Harrimaniidae | 3 |
| Mollusca;Bivalvia;Heterodonta;Euheterodonta;Lucinoida;Thyasiroidea;Thyasiridae | 1 |
| Mollusca;Gastropoda;Heterobranchia;Euthyneura;...Cavoliniidae | 1 |
| Nemertea;Enopla;Hoplonemertea;Monostilifera;Eumonostilifera;Amphiporidae | 1 |
| Platyhelminthes;Rhabditophora;Seriata;Tricladida;Maricola;Procerodoidea;Procerodidae | 1 |
| Porifera;Calcarea;Calcaronea;Leucosolenida;Grantiidae | 1 |
| Porifera;Calcarea;Calcinea;Clathrinida;Soleneiscidae | 1 |

**Table S5.** Reference sequences used in the hydrozoan phylogenetic trees (Figs. S5).

| Accession Code | Simplified taxonomic lineage |
|---|---|
| **EU190843** | Anthozoa Hexacorallia Actinostola crassicornis |
| **EF589071** | Anthozoa Hexacorallia Amplexidiscus fenestrafer |
| **X53498** | Anthozoa Hexacorallia Anemonia sulcata |
| **Z21671** | Anthozoa Hexacorallia Anthopleura kurogane |
| **Z92908** | Anthozoa Hexacorallia Antipathes lata |
| **EU190852** | Anthozoa Hexacorallia Boloceroides mcmurrichi |
| **EF589065** | Anthozoa Hexacorallia Corynactis viridis |
| **Z86097** | Anthozoa Hexacorallia Diadumene lineata |
| **Z92904** | Anthozoa Hexacorallia Epiactis japonica |
| **Z92905** | Anthozoa Hexacorallia Flosmaris mutsuensis |
| **EF589074** | Anthozoa Hexacorallia Fungiacyathus marenzelleri |
| **U42453** | Anthozoa Hexacorallia Parazoanthus axinellae |
| **EU190871** | Anthozoa Hexacorallia Phymanthus loligo |
| **LT631265** | Anthozoa Hexacorallia Pseudodiploria clivosa |
| **Z92907** | Anthozoa Hexacorallia Rhizopsammia minuta |
| **EF589072** | Anthozoa Hexacorallia Rhodactis rhodostoma |
| **Z92906** | Anthozoa Hexacorallia Tubastraea coccinea |
| **Z49195** | Anthozoa Octocorallia Bellonella rigida |
| **Z92900** | Anthozoa Octocorallia Calicogorgia granulosa |
| **KP324638** | Anthozoa Octocorallia Dasystenella austasensis |
| **KP324679** | Anthozoa Octocorallia Fanellia korema |
| **KP324675** | Anthozoa Octocorallia Fanellia medialis |
| **KP324648** | Anthozoa Octocorallia Fannyella rossii |
| **Z92903** | Anthozoa Octocorallia Leioptilus fimbriatus |
| **KP324716** | Anthozoa Octocorallia Metafannyella kuekenthali |
| **KP324636** | Anthozoa Octocorallia Onogorgia nodosa |
| **KP324695** | Anthozoa Octocorallia Parastenella spinosa |
| **KP324674** | Anthozoa Octocorallia Perissogorgia vitrea |
| **KP324702** | Anthozoa Octocorallia Primnoella antarctica |
| **KP324736** | Anthozoa Octocorallia Primnoella chilensis |
| **KP324715** | Anthozoa Octocorallia Primnoella divaricata |
| **Z86106** | Anthozoa Octocorallia Virgularia gustaviana |
| **MF599314** | Ctenophora Lobatolampea tetragona |

| | |
|---|---|
| **LC047805** | Cubozoa Alatina philippina |
| **AF358107** | Cubozoa Carukia barnesi |
| **AF358108** | Cubozoa Carybdea rastonii |
| **AF358104** | Cubozoa Chironex fleckeri |
| **AF358103** | Cubozoa Chiropsalmus AGC-2001 |
| **GQ849084** | Cubozoa Malo kingi |
| **LC033479** | Cubozoa Meteorona kishinouyei |
| **GQ849086** | Cubozoa Tamoya ohboya |
| **EU876578** | Hydrozoa Hydroidolina Anthoathecata Capitata Asyncoryne ryniensis |
| **EU876578** | Hydrozoa Hydroidolina Anthoathecata Capitata Asyncoryne ryniensis |
| **LT593888** | Hydrozoa Hydroidolina Anthoathecata Capitata Cladocorynidae Pteroclava krempfi |
| **GQ424329** | Hydrozoa Hydroidolina Anthoathecata Capitata Corynidae Coryne nipponica |
| **GQ424329** | Hydrozoa Hydroidolina Anthoathecata Capitata Corynidae Coryne nipponica |
| **EU883544** | Hydrozoa Hydroidolina Anthoathecata Capitata Corynidae Dipurena halterata |
| **EU883544** | Hydrozoa Hydroidolina Anthoathecata Capitata Corynidae Dipurena halterata |
| **GQ424334** | Hydrozoa Hydroidolina Anthoathecata Capitata Corynidae Dipurena reesi |
| **GQ424334** | Hydrozoa Hydroidolina Anthoathecata Capitata Corynidae Dipurena reesi |
| **GQ424337** | Hydrozoa Hydroidolina Anthoathecata Capitata Corynidae Sarsia lovenii |
| **GQ424337** | Hydrozoa Hydroidolina Anthoathecata Capitata Corynidae Sarsia lovenii |
| **GQ424339** | Hydrozoa Hydroidolina Anthoathecata Capitata Corynidae Sarsia tubulosa |
| **GQ424339** | Hydrozoa Hydroidolina Anthoathecata Capitata Corynidae Sarsia tubulosa |
| **LT631288** | Hydrozoa Hydroidolina Anthoathecata Capitata Milleporidae Millepora GA112 |
| **LT631287** | Hydrozoa Hydroidolina Anthoathecata Capitata Milleporidae Millepora DJ207 |
| **LT631288** | Hydrozoa Hydroidolina Anthoathecata Capitata Milleporidae Millepora GA112 |
| **LT631287** | Hydrozoa Hydroidolina Anthoathecata Capitata Milleporidae Millepora DJ207 |
| **GQ424344** | Hydrozoa Hydroidolina Anthoathecata Capitata Polyorchidae Polyorchis haplus |
| **GQ424344** | Hydrozoa Hydroidolina Anthoathecata Capitata Polyorchidae Polyorchis haplus |
| **KF962297** | Hydrozoa Hydroidolina Anthoathecata FileferaIV Turritopsis lata |
| **JQ407395** | Hydrozoa Hydroidolina Anthoathecata Filifera Bouillonactinia hooperi |
| **JQ407395** | Hydrozoa Hydroidolina Anthoathecata Filifera Bouillonactinia hooperi |
| **JQ407378** | Hydrozoa Hydroidolina Anthoathecata Filifera Hydractinia echinata |
| **JQ407378** | Hydrozoa Hydroidolina Anthoathecata Filifera Hydractinia echinata |
| **EU272625** | Hydrozoa Hydroidolina Anthoathecata Filifera Lizzia blondina |
| **EU448097** | Hydrozoa Hydroidolina Anthoathecata Filifera Neoturris breviconis |
| **JQ407388** | Hydrozoa Hydroidolina Anthoathecata Filifera Podocoryna borealis |
| **JQ407388** | Hydrozoa Hydroidolina Anthoathecata Filifera Podocoryna borealis |

| | |
|---|---|
| **EU272634** | Hydrozoa Hydroidolina Anthoathecata Filifera Rathkea octopunctata |
| **JQ407403** | Hydrozoa Hydroidolina Anthoathecata Filifera Schuchertinia milleri |
| **JQ407403** | Hydrozoa Hydroidolina Anthoathecata Filifera Schuchertinia milleri |
| **KT722442** | Hydrozoa Hydroidolina Anthoathecata Filifera Turritopsis nutricula |
| **AY920767** | Hydrozoa Hydroidolina Anthoathecata FiliferaII Fabienna sphaerica |
| **EU272622** | Hydrozoa Hydroidolina Anthoathecata FiliferaII Hydrichthella epigorgia |
| **EU272622** | Hydrozoa Hydroidolina Anthoathecata FiliferaII Hydrichthella epigorgia |
| **EU305500** | Hydrozoa Hydroidolina Anthoathecata FiliferaII Proboscidactyla flavicirrata |
| **EU272631** | Hydrozoa Hydroidolina Anthoathecata FiliferaII Proboscidactyla ornata |
| **JQ407377** | Hydrozoa Hydroidolina Anthoathecata FiliferaIII Hydractinia symbiolongicarpus |
| **JQ407377** | Hydrozoa Hydroidolina Anthoathecata FiliferaIII Hydractinia symbiolongicarpus |
| **JQ407367** | Hydrozoa Hydroidolina Anthoathecata FiliferaIII Hydractiniidae Clava multicornis |
| **JQ407367** | Hydrozoa Hydroidolina Anthoathecata FiliferaIII Hydractiniidae Clava multicornis |
| **JQ407363** | Hydrozoa Hydroidolina Anthoathecata FiliferaIII Janaria mirabilis |
| **JQ407363** | Hydrozoa Hydroidolina Anthoathecata FiliferaIII Janaria mirabilis |
| **EU272644** | Hydrozoa Hydroidolina Anthoathecata FiliferaIII Lepidopora microstylus |
| **EU272644** | Hydrozoa Hydroidolina Anthoathecata FiliferaIII Lepidopora microstylus |
| **AF358092** | Hydrozoa Hydroidolina Anthoathecata FiliferaIII Podocoryna exigua |
| **AF358092** | Hydrozoa Hydroidolina Anthoathecata FiliferaIII Podocoryna exigua |
| **KT722386** | Hydrozoa Hydroidolina Anthoathecata FiliferaIV Bimeria vestita |
| **KT722386** | Hydrozoa Hydroidolina Anthoathecata FiliferaIV Bimeria vestita |
| **MG788989** | Hydrozoa Hydroidolina Anthoathecata FiliferaIV Bougainvillia carolinensis |
| **EU272606** | Hydrozoa Hydroidolina Anthoathecata FiliferaIV Bougainvillia carolinensis |
| **MG788989** | Hydrozoa Hydroidolina Anthoathecata FiliferaIV Bougainvillia carolinensis |
| **EU272606** | Hydrozoa Hydroidolina Anthoathecata FiliferaIV Bougainvillia carolinensis |
| **EU305490** | Hydrozoa Hydroidolina Anthoathecata FiliferaIV Bougainvillia fulva |
| **EU305490** | Hydrozoa Hydroidolina Anthoathecata FiliferaIV Bougainvillia fulva |
| **FJ550582** | Hydrozoa Hydroidolina Anthoathecata FiliferaIV Bougainvillia muscus |
| **FJ550582** | Hydrozoa Hydroidolina Anthoathecata FiliferaIV Bougainvillia muscus |
| **EU272632** | Hydrozoa Hydroidolina Anthoathecata FiliferaIV Garveia grisea |
| **EU272623** | Hydrozoa Hydroidolina Anthoathecata FiliferaIV Koellikerina fasciculata |
| **EU272623** | Hydrozoa Hydroidolina Anthoathecata FiliferaIV Koellikerina fasciculata |
| **EU272625** | Hydrozoa Hydroidolina Anthoathecata FiliferaIV Lizzia blondina |
| **EU272627** | Hydrozoa Hydroidolina Anthoathecata FiliferaIV Pachycordyle pusilla |
| **EU272634** | Hydrozoa Hydroidolina Anthoathecata FiliferaIV Rathkea octopunctata |
| **EU272635** | Hydrozoa Hydroidolina Anthoathecata FiliferaIV Rhizogeton nudus |

| | |
|---|---|
| **KT722442** | Hydrozoa Hydroidolina Anthoathecata FiliferaIV Turritopsis nutricula |
| **EU876564** | Hydrozoa Hydroidolina Aplanulata Corymorpha bigelowi |
| **EU876564** | Hydrozoa Hydroidolina Aplanulata Corymorpha bigelowi |
| **EU876565** | Hydrozoa Hydroidolina Aplanulata Corymorpha pendula |
| **EU876565** | Hydrozoa Hydroidolina Aplanulata Corymorpha pendula |
| **EU883547** | Hydrozoa Hydroidolina Aplanulata Ectopleura marina |
| **EU883547** | Hydrozoa Hydroidolina Aplanulata Ectopleura marina |
| **EU876566** | Hydrozoa Hydroidolina Aplanulata Hybocodon chilensis |
| **EU876566** | Hydrozoa Hydroidolina Aplanulata Hybocodon chilensis |
| **FJ265733** | Hydrozoa Hydroidolina Aplanulata Hydra sinensis |
| **FJ265733** | Hydrozoa Hydroidolina Aplanulata Hydra sinensis |
| **KF962297** | Hydrozoa Hydroidolina FileferaIV Turritopsis lata |
| **KT722442** | Hydrozoa Hydroidolina FiliferaIV Turritopsis nutricula |
| **AF358076** | Hydrozoa Hydroidolina Leptothecata Aequoreidae Aequorea forskalea |
| **AF358076** | Hydrozoa Hydroidolina Leptothecata Aequoreidae Aequorea forskalea |
| **EU305501** | Hydrozoa Hydroidolina Leptothecata Aequoreidae Rhacostoma atlanticum |
| **EU305501** | Hydrozoa Hydroidolina Leptothecata Aequoreidae Rhacostoma atlanticum |
| **EU272601** | Hydrozoa Hydroidolina Leptothecata Aglaopheniidae Aglaophenia tubiformis |
| **EU272601** | Hydrozoa Hydroidolina Leptothecata Aglaopheniidae Aglaophenia tubiformis |
| **AY789739** | Hydrozoa Hydroidolina Leptothecata Campanulariidae Campanularia volubilis |
| **AY789739** | Hydrozoa Hydroidolina Leptothecata Campanulariidae Campanularia volubilis |
| **AY789737** | Hydrozoa Hydroidolina Leptothecata Campanulariidae Orthopyxis integra |
| **AY789737** | Hydrozoa Hydroidolina Leptothecata Campanulariidae Orthopyxis integra |
| **AY789738** | Hydrozoa Hydroidolina Leptothecata Campanulariidae Rhizocaulus verticillatus |
| **AY789738** | Hydrozoa Hydroidolina Leptothecata Campanulariidae Rhizocaulus verticillatus |
| **FJ550549** | Hydrozoa Hydroidolina Leptothecata Campanulariidae Silicularia rosea |
| **FJ550549** | Hydrozoa Hydroidolina Leptothecata Campanulariidae Silicularia rosea |
| **KF962267** | Hydrozoa Hydroidolina Leptothecata Eirenidae Eirene kambara |
| **KF962267** | Hydrozoa Hydroidolina Leptothecata Eirenidae Eirene kambara |
| **FJ550588** | Hydrozoa Hydroidolina Leptothecata Eirenidae Eirene viridula |
| **FJ550588** | Hydrozoa Hydroidolina Leptothecata Eirenidae Eirene viridula |
| **FJ550599** | Hydrozoa Hydroidolina Leptothecata Eirenidae Eutima curva |
| **FJ550599** | Hydrozoa Hydroidolina Leptothecata Eirenidae Eutima curva |
| **FJ550600** | Hydrozoa Hydroidolina Leptothecata Eirenidae Eutima gegenbauri |
| **FJ550600** | Hydrozoa Hydroidolina Leptothecata Eirenidae Eutima gegenbauri |
| **KY363986** | Hydrozoa Hydroidolina Leptothecata Eirenidae Eutima gracilis |

| | |
|---|---|
| **KY363986** | Hydrozoa Hydroidolina Leptothecata Eirenidae Eutima gracilis |
| **EU305493** | Hydrozoa Hydroidolina Leptothecata Eirenidae Eutima sapinhoa |
| **EU305493** | Hydrozoa Hydroidolina Leptothecata Eirenidae Eutima sapinhoa |
| **KY363971** | Hydrozoa Hydroidolina Leptothecata Eirenidae Eutonina indicans |
| **KY363971** | Hydrozoa Hydroidolina Leptothecata Eirenidae Eutonina indicans |
| **KY363989** | Hydrozoa Hydroidolina Leptothecata Eirenidae Helgicirrha cari |
| **KY363989** | Hydrozoa Hydroidolina Leptothecata Eirenidae Helgicirrha cari |
| **KY363983** | Hydrozoa Hydroidolina Leptothecata Laodiceidae Ptychogena crocea |
| **KY363983** | Hydrozoa Hydroidolina Leptothecata Laodiceidae Ptychogena crocea |
| **FJ550535** | Hydrozoa Hydroidolina Leptothecata Laodiceidae Staurodiscus gotoi |
| **KY363978** | Hydrozoa Hydroidolina Leptothecata Laodiceidae Staurostoma mertensii |
| **KY363978** | Hydrozoa Hydroidolina Leptothecata Laodiceidae Staurostoma mertensii |
| **KY363984** | Hydrozoa Hydroidolina Leptothecata Mitrocomidae Cyclocanna welshi |
| **KY363984** | Hydrozoa Hydroidolina Leptothecata Mitrocomidae Cyclocanna welshi |
| **KY363982** | Hydrozoa Hydroidolina Leptothecata Mitrocomidae Earleria quadrata |
| **KY363982** | Hydrozoa Hydroidolina Leptothecata Mitrocomidae Earleria quadrata |
| **KY363977** | Hydrozoa Hydroidolina Leptothecata Mitrocomidae Halopsis ocellata |
| **KY363977** | Hydrozoa Hydroidolina Leptothecata Mitrocomidae Halopsis ocellata |
| **FJ550536** | Hydrozoa Hydroidolina Leptothecata Mitrocomidae Mitrocomella niwai |
| **FJ550536** | Hydrozoa Hydroidolina Leptothecata Mitrocomidae Mitrocomella niwai |
| **KY363979** | Hydrozoa Hydroidolina Leptothecata Mitrocomidae Mitrocomella polydiademata |
| **KY363979** | Hydrozoa Hydroidolina Leptothecata Mitrocomidae Mitrocomella polydiademata |
| **KF962217** | Hydrozoa Hydroidolina Leptothecata Obeliida Clytiidae Clytia folleata |
| **KF962217** | Hydrozoa Hydroidolina Leptothecata Obeliida Clytiidae Clytia folleata |
| **KX665388** | Hydrozoa Hydroidolina Leptothecata Obeliida Clytiidae Clytia gracilis |
| **KX665388** | Hydrozoa Hydroidolina Leptothecata Obeliida Clytiidae Clytia gracilis |
| **KX665400** | Hydrozoa Hydroidolina Leptothecata Obeliida Clytiidae Clytia hemisphaerica |
| **KX665400** | Hydrozoa Hydroidolina Leptothecata Obeliida Clytiidae Clytia hemisphaerica |
| **KX665454** | Hydrozoa Hydroidolina Leptothecata Obeliida Clytiidae Clytia noliformis |
| **KX665454** | Hydrozoa Hydroidolina Leptothecata Obeliida Clytiidae Clytia noliformis |
| **KX665404** | Hydrozoa Hydroidolina Leptothecata Obeliida Obeliidae Laomedea angulata |
| **KX665404** | Hydrozoa Hydroidolina Leptothecata Obeliida Obeliidae Laomedea angulata |
| **KX665446** | Hydrozoa Hydroidolina Leptothecata Obeliida Obeliidae Laomedea calceolifera |
| **KX665446** | Hydrozoa Hydroidolina Leptothecata Obeliida Obeliidae Laomedea calceolifera |
| **KX665412** | Hydrozoa Hydroidolina Leptothecata Obeliida Obeliidae Obelia bidentata |
| **KX665412** | Hydrozoa Hydroidolina Leptothecata Obeliida Obeliidae Obelia bidentata |

| | |
|---|---|
| **MG792325** | Hydrozoa Hydroidolina Leptothecata Obeliida Obeliidae Obelia dichotoma |
| **MG792325** | Hydrozoa Hydroidolina Leptothecata Obeliida Obeliidae Obelia dichotoma |
| **KT722412** | Hydrozoa Hydroidolina Leptothecata Obeliida Obeliidae Obelia geniculata |
| **KT722412** | Hydrozoa Hydroidolina Leptothecata Obeliida Obeliidae Obelia geniculata |
| **EU305499** | Hydrozoa Hydroidolina Leptothecata Plumulariidae Plumularia hyalina |
| **EU305499** | Hydrozoa Hydroidolina Leptothecata Plumulariidae Plumularia hyalina |
| **KT722433** | Hydrozoa Hydroidolina Leptothecata Sertulariidae Sertularella cylindritheca |
| **KT722433** | Hydrozoa Hydroidolina Leptothecata Sertulariidae Sertularella cylindritheca |
| **FJ550540** | Hydrozoa Hydroidolina Leptothecata Tiarannidae Modeeria rotunda |
| **FJ550540** | Hydrozoa Hydroidolina Leptothecata Tiarannidae Modeeria rotunda |
| **FJ550598** | Hydrozoa Hydroidolina Leptothecata Tiarannidae Stegopoma plicatile |
| **FJ550598** | Hydrozoa Hydroidolina Leptothecata Tiarannidae Stegopoma plicatile |
| **AF358079** | Hydrozoa Hydroidolina Leptothecata Tiaropsidae Tiaropsidium kelseyi |
| **AF358079** | Hydrozoa Hydroidolina Leptothecata Tiaropsidae Tiaropsidium kelseyi |
| **FJ550531** | Hydrozoa Hydroidolina Leptothecata Tiaropsidae Tiaropsis multicirrata |
| **FJ550531** | Hydrozoa Hydroidolina Leptothecata Tiaropsidae Tiaropsis multicirrata |
| **AY937345** | Hydrozoa Hydroidolina Siphonophorae Calycophorae Abylopsis tetragona |
| **AY937345** | Hydrozoa Hydroidolina Siphonophorae Calycophorae Abylopsis tetragona |
| **KX421847** | Hydrozoa Hydroidolina Siphonophorae Calycophorae Chuniphyes moserae |
| **KX421847** | Hydrozoa Hydroidolina Siphonophorae Calycophorae Chuniphyes moserae |
| **AY937335** | Hydrozoa Hydroidolina Siphonophorae Calycophorae Chuniphyes multidentata |
| **AY937335** | Hydrozoa Hydroidolina Siphonophorae Calycophorae Chuniphyes multidentata |
| **KX421848** | Hydrozoa Hydroidolina Siphonophorae Calycophorae Clausophyes moserae |
| **KX421848** | Hydrozoa Hydroidolina Siphonophorae Calycophorae Clausophyes moserae |
| **AY937336** | Hydrozoa Hydroidolina Siphonophorae Calycophorae Clausophyes ovata |
| **AY937336** | Hydrozoa Hydroidolina Siphonophorae Calycophorae Clausophyes ovata |
| **KX421850** | Hydrozoa Hydroidolina Siphonophorae Calycophorae Crystallophyes amygdalina |
| **KX421850** | Hydrozoa Hydroidolina Siphonophorae Calycophorae Crystallophyes amygdalina |
| **AY937318** | Hydrozoa Hydroidolina Siphonophorae Calycophorae Diphyes dispar |
| **AY937318** | Hydrozoa Hydroidolina Siphonophorae Calycophorae Diphyes dispar |
| **DQ080014** | Hydrozoa Hydroidolina Siphonophorae Calycophorae Lilyopsis rosea |
| **DQ080014** | Hydrozoa Hydroidolina Siphonophorae Calycophorae Lilyopsis rosea |
| **AY937337** | Hydrozoa Hydroidolina Siphonophorae Calycophorae Muggiaea atlantica |
| **AY937337** | Hydrozoa Hydroidolina Siphonophorae Calycophorae Muggiaea atlantica |
| **KX421851** | Hydrozoa Hydroidolina Siphonophorae Calycophorae Prayidae Desmophyes haematogaster |
| **KX421851** | Hydrozoa Hydroidolina Siphonophorae Calycophorae Prayidae Desmophyes haematogaster |

| | |
|---|---|
| **AY919607** | Hydrozoa Hydroidolina Siphonophorae Calycophorae Prayidae Lilyopsis fluoracantha |
| **AY919607** | Hydrozoa Hydroidolina Siphonophorae Calycophorae Prayidae Lilyopsis fluoracantha |
| **KX421854** | Hydrozoa Hydroidolina Siphonophorae Calycophorae Sphaeronectes haddocki |
| **KX421854** | Hydrozoa Hydroidolina Siphonophorae Calycophorae Sphaeronectes haddocki |
| **AF358065** | Hydrozoa Hydroidolina Siphonophorae Cystonectae Physalia physalis |
| **AF358065** | Hydrozoa Hydroidolina Siphonophorae Cystonectae Physalia physalis |
| **AY937327** | Hydrozoa Hydroidolina Siphonophorae Cystonectae Rhizophysa filiformis |
| **AY937327** | Hydrozoa Hydroidolina Siphonophorae Cystonectae Rhizophysa filiformis |
| **AY937312** | Hydrozoa Hydroidolina Siphonophorae Physonectae Agalma clausi |
| **AY937312** | Hydrozoa Hydroidolina Siphonophorae Physonectae Agalma clausi |
| **KF214714** | Hydrozoa Hydroidolina Siphonophorae Physonectae Apolemia lanosa |
| **KF214714** | Hydrozoa Hydroidolina Siphonophorae Physonectae Apolemia lanosa |
| **KF214715** | Hydrozoa Hydroidolina Siphonophorae Physonectae Apolemia rubriversa |
| **KF214715** | Hydrozoa Hydroidolina Siphonophorae Physonectae Apolemia rubriversa |
| **AY937334** | Hydrozoa Hydroidolina Siphonophorae Physonectae Bargmannia elongata |
| **AY937334** | Hydrozoa Hydroidolina Siphonophorae Physonectae Bargmannia elongata |
| **AY937317** | Hydrozoa Hydroidolina Siphonophorae Physonectae Cordagalma cordiforme |
| **AY937317** | Hydrozoa Hydroidolina Siphonophorae Physonectae Cordagalma cordiforme |
| **AY937319** | Hydrozoa Hydroidolina Siphonophorae Physonectae Forskalia asymmetrica |
| **AY937319** | Hydrozoa Hydroidolina Siphonophorae Physonectae Forskalia asymmetrica |
| **EU301622** | Hydrozoa Trachylinae Actinulida Halammohydra BB-2008 |
| **EU301622** | Hydrozoa Trachylinae Actinulida Halammohydra BB-2008 |
| **EU301622** | Hydrozoa Trachylinae Actinulida Halammohydra USNM 1109974 |
| **EU301622** | Hydrozoa Trachylinae Actinulida Halammohydra USNM 1109974 |
| **KY077286** | Hydrozoa Trachylinae Limnomedusae Astrohydra japonica |
| **KY077286** | Hydrozoa Trachylinae Limnomedusae Astrohydra japonica |
| **KF771643** | Hydrozoa Trachylinae Limnomedusae Craspedacusta sowerbii |
| **KF771643** | Hydrozoa Trachylinae Limnomedusae Craspedacusta sowerbii |
| **AY920755** | Hydrozoa Trachylinae Limnomedusae Limnocnida tanganjicae |
| **AY920755** | Hydrozoa Trachylinae Limnomedusae Limnocnida tanganjicae |
| **AF358056** | Hydrozoa Trachylinae Limnomedusae Maeotias marginata |
| **AF358056** | Hydrozoa Trachylinae Limnomedusae Maeotias marginata |
| **EU247813** | Hydrozoa Trachylinae Narcomedusae Aeginidae Aegina citrea |
| **EU247813** | Hydrozoa Trachylinae Narcomedusae Aeginidae Aegina citrea |
| **MG979347** | Hydrozoa Trachylinae Narcomedusae Aeginidae Aeginopsis laurentii |
| **MG979347** | Hydrozoa Trachylinae Narcomedusae Aeginidae Aeginopsis laurentii |

| | |
|---|---|
| **KY007605** | Hydrozoa Trachylinae Narcomedusae Aeginidae Aeginura grimaldii |
| **KY007605** | Hydrozoa Trachylinae Narcomedusae Aeginidae Aeginura grimaldii |
| **MG979341** | Hydrozoa Trachylinae Narcomedusae Aeginidae Bathykorus bouilloni |
| **MG979341** | Hydrozoa Trachylinae Narcomedusae Aeginidae Bathykorus bouilloni |
| **EU247812** | Hydrozoa Trachylinae Narcomedusae Aeginidae Solmundella bitentaculata |
| **EU247812** | Hydrozoa Trachylinae Narcomedusae Aeginidae Solmundella bitentaculata |
| **AF358059** | Hydrozoa Trachylinae Narcomedusae Cuninidae Cunina frugifera |
| **AF358059** | Hydrozoa Trachylinae Narcomedusae Cuninidae Cunina frugifera |
| **KY007606** | Hydrozoa Trachylinae Narcomedusae Cuninidae Cunina octonaria |
| **KY007606** | Hydrozoa Trachylinae Narcomedusae Cuninidae Cunina octonaria |
| **KY007607** | Hydrozoa Trachylinae Narcomedusae Cuninidae Sigiweddellia sp BB-2016 |
| **KY007607** | Hydrozoa Trachylinae Narcomedusae Cuninidae Sigiweddellia sp BB-2016 |
| **MG979342** | Hydrozoa Trachylinae Narcomedusae Cuninidae Solmissus albescens |
| **MG979342** | Hydrozoa Trachylinae Narcomedusae Cuninidae Solmissus albescens |
| **AF358060** | Hydrozoa Trachylinae Narcomedusae Cuninidae Solmissus marshalli |
| **AF358060** | Hydrozoa Trachylinae Narcomedusae Cuninidae Solmissus marshalli |
| **MG979344** | Hydrozoa Trachylinae Narcomedusae Solmarisidae Pegantha martagon |
| **MG979344** | Hydrozoa Trachylinae Narcomedusae Solmarisidae Pegantha martagon |
| **MG979345** | Hydrozoa Trachylinae Narcomedusae Solmarisidae Pegantha rubiginosa |
| **MG979345** | Hydrozoa Trachylinae Narcomedusae Solmarisidae Pegantha rubiginosa |
| **MG979346** | Hydrozoa Trachylinae Narcomedusae Solmarisidae Solmaris DLSI359 |
| **MG979346** | Hydrozoa Trachylinae Narcomedusae Solmarisidae Solmaris DLSI359 |
| **MG979348** | Hydrozoa Trachylinae Narcomedusae Tetraplatiidae Tetraplatia chuni |
| **MG979348** | Hydrozoa Trachylinae Narcomedusae Tetraplatiidae Tetraplatia chuni |
| **DQ002501** | Hydrozoa Trachylinae Narcomedusae Tetraplatiidae Tetraplatia volitans |
| **DQ002501** | Hydrozoa Trachylinae Narcomedusae Tetraplatiidae Tetraplatia volitans |
| **EU247816** | Hydrozoa Trachylinae Trachymedusae Geryoniidae Geryonia proboscidalis |
| **EU247816** | Hydrozoa Trachylinae Trachymedusae Geryoniidae Geryonia proboscidalis |
| **AY920756** | Hydrozoa Trachylinae Trachymedusae Geryoniidae Liriope tetraphylla |
| **AY920756** | Hydrozoa Trachylinae Trachymedusae Geryoniidae Liriope tetraphylla |
| **EU247822** | Hydrozoa Trachylinae Trachymedusae Halicreatidae Botrynema brucei |
| **EU247826** | Hydrozoa Trachylinae Trachymedusae Halicreatidae Halicreas minimum |
| **EU247825** | Hydrozoa Trachylinae Trachymedusae Halicreatidae Haliscera conica |
| **MG979352** | Hydrozoa Trachylinae Trachymedusae Halicreatidae Halitrephes maasi |
| **EU247821** | Hydrozoa Trachylinae Trachymedusae Rhopalonematidae Aglantha digitale |
| **EU247821** | Hydrozoa Trachylinae Trachymedusae Rhopalonematidae Aglantha digitale |

| | |
|---|---|
| **EU247820** | Hydrozoa Trachylinae Trachymedusae Rhopalonematidae Aglaura hemistoma |
| **EU247820** | Hydrozoa Trachylinae Trachymedusae Rhopalonematidae Aglaura hemistoma |
| **MG979355** | Hydrozoa Trachylinae Trachymedusae Rhopalonematidae Amphogona apicata |
| **MG979355** | Hydrozoa Trachylinae Trachymedusae Rhopalonematidae Amphogona apicata |
| **MK547161** | Hydrozoa Trachylinae Trachymedusae Rhopalonematidae Benthocodon peduncula |
| **MK547161** | Hydrozoa Trachylinae Trachymedusae Rhopalonematidae Benthocodon peduncula |
| **MG979358** | Hydrozoa Trachylinae Trachymedusae Rhopalonematidae Colobonema sericeum |
| **MG979358** | Hydrozoa Trachylinae Trachymedusae Rhopalonematidae Colobonema sericeum |
| **MK547162** | Hydrozoa Trachylinae Trachymedusae Rhopalonematidae Crossota alba |
| **MK547162** | Hydrozoa Trachylinae Trachymedusae Rhopalonematidae Crossota alba |
| **MG979362** | Hydrozoa Trachylinae Trachymedusae Rhopalonematidae Crossota norvegica |
| **MG979362** | Hydrozoa Trachylinae Trachymedusae Rhopalonematidae Crossota norvegica |
| **AF358063** | Hydrozoa Trachylinae Trachymedusae Rhopalonematidae Crossota rufobrunnea |
| **AF358063** | Hydrozoa Trachylinae Trachymedusae Rhopalonematidae Crossota rufobrunnea |
| **AF358062** | Hydrozoa Trachylinae Trachymedusae Rhopalonematidae Pantachogon haeckeli |
| **AF358062** | Hydrozoa Trachylinae Trachymedusae Rhopalonematidae Pantachogon haeckeli |
| **EU247819** | Hydrozoa Trachylinae Trachymedusae Rhopalonematidae Rhopalonema velatum |
| **EU247819** | Hydrozoa Trachylinae Trachymedusae Rhopalonematidae Rhopalonema velatum |
| **MG979364** | Hydrozoa Trachylinae Trachymedusae Rhopalonematidae Sminthea arctica |
| **MG979364** | Hydrozoa Trachylinae Trachymedusae Rhopalonematidae Sminthea arctica |
| **KY077285** | Hydrozoa Trachylinae Trachymedusae Tetrorchis erythrogaster |
| **KY077285** | Hydrozoa Trachylinae Trachymedusae Tetrorchis erythrogaster |
| **AY920771** | Scyphozoa Cassiopea xamachana |
| **AF358100** | Scyphozoa Catostylus AGC-2001 |
| **KY610843** | Scyphozoa Chrysaora chinensis |
| **AF358099** | Scyphozoa Chrysaora melanaster |
| **KY212123** | Scyphozoa Chrysaora pacifica |
| **JX845355** | Scyphozoa Cyanea nozakii |
| **HM194791** | Scyphozoa Deepstaria enigmatica |
| **KY610748** | Scyphozoa Lobonema smithii |
| **AF358096** | Scyphozoa Phacellophora camtschatica |
| **HM194792** | Scyphozoa Poralia rufescens |
| **AY920772** | Scyphozoa Rhizostome sp. AGC-2005 |
| **JX845352** | Scyphozoa Rhopilema esculentum |
| **AF358101** | Scyphozoa Stomolophus meleagris |
| **KU308576** | Staurozoa Calvadosia cruciformis |

| | |
|---|---|
| **KU308571** | Staurozoa Calvadosia cruxmelitensis |
| **KU308558** | Staurozoa Depastromorpha africana |
| **AF358102** | Staurozoa Haliclystus sanjuanensis |
| **AY845345** | Staurozoa Lucernaria janetae |

**Table S6.** Reference sequences used in the phylogenetic tree of Copepoda (Fig. S6).

| Accession Code | Simplified taxonomic lineage |
|---|---|
| **JX995284** | Calanoida Acartia clausii |
| **GU969197** | Calanoida Acartia danae |
| **KY859764** | Calanoida Acartia hudsonica |
| **GU969198** | Calanoida Acartia negligens |
| **GU969196** | Calanoida Acartia omorii |
| **GU969157** | Calanoida Acartia pacifica |
| **KT030253** | Calanoida Acartia steueri |
| **GU350740** | Calanoida Acartia tonsa |
| **MG660981** | Calanoida Aetideidae Chiridiella reductella |
| **MG660938** | Calanoida Aetideidae Chiridius obtusifrons |
| **AB625971** | Calanoida Aetideidae Euchirella amoena |
| **MG660908** | Calanoida Aetideidae Gaetanus brevispinus |
| **MG660879** | Calanoida Aetideidae Gaetanus tenuispinus |
| **MG661009** | Calanoida Aetideidae Jaschnovia brevis |
| **MF796518** | Calanoida Aetideidae Prolutamator hadalis |
| **MG660990** | Calanoida Aetideidae Pseudochirella batillipa |
| **AB625972** | Calanoida Aetideidae Undeuchaeta major |
| **HM997058** | Calanoida Arietellidae Paraugaptilus buchani |
| **MG660988** | Calanoida Augaptilidae Haloptilus acutifrons |
| **GU969152** | Calanoida Augaptilidae Haloptilus longicornis |
| **MG660876** | Calanoida Augaptilidae Pseudaugaptilus polaris |
| **MG661046** | Calanoida Augaptilidae Pseudhaloptilus pacificus |
| **MG660956** | Calanoida Bathypontiidae Temorites brevis |
| **MG660955** | Calanoida Bathypontiidae Temorites brevis |
| **JQ911941** | Calanoida Bestiolina similis |
| **MF993124** | Calanoida Calanidae Calanus Calanidae Calanus finmarchicus |
| **MF993123** | Calanoida Calanidae Calanus Calanidae Calanus glacialis |
| **AY118066** | Calanoida Calanidae Calanus Calanidae Calanus propinquus |
| **AF367716** | Calanoida Calanidae Mesocalanus tenuicornis |
| **JQ911943** | Calanoida Calocalanidae Calocalanus curtus |
| **GU969160** | Calanoida Calocalanidae Calocalanus pavo |
| **GU969146** | Calanoida Calocalanidae Calocalanus plumulosus |
| **AB625973** | Calanoida Candacia bipinnata |

| | |
|---|---|
| **GU969213** | Calanoida Candacia bispinosa |
| **AB625974** | Calanoida Candacia columbiae |
| **GU969145** | Calanoida Candacia discaudata |
| **GU969199** | Calanoida Candacia pachydactyla |
| **GU969161** | Calanoida Candacia truncata |
| **GU969163** | Calanoida Centropagidae Centropages abdominalis |
| **GU969158** | Calanoida Centropagidae Centropages furcatus |
| **JX995295** | Calanoida Centropagidae Centropages hamatus |
| **GU969162** | Calanoida Centropagidae Centropages tenuiremis |
| **JX995298** | Calanoida Centropagidae Centropages typicus |
| **JX995304** | Calanoida Centropagidae Isias clavipes |
| **GU969144** | Calanoida Centropagidae Sinocalanus tenellus |
| **GU969200** | Calanoida Clausocalanidae Clausocalanus furcatus |
| **JX995321** | Calanoida Clausocalanidae Pseudocalanus elongatus |
| **JX995324** | Calanoida Clausocalanidae Pseudocalanus moultoni |
| **GU969206** | Calanoida Cosmocalanus darwinii |
| **AY339147** | Calanoida Diaptomidae Arctodiaptomus dorsalis |
| **AY339149** | Calanoida Diaptomidae Eudiaptomus graciloides |
| **JX945140** | Calanoida Diaptomidae Hemidiaptomus roubaui |
| **AY339155** | Calanoida Diaptomidae Leptodiaptomus sicilis |
| **AY339156** | Calanoida Diaptomidae Mastigodiaptomus nesus |
| **AY339159** | Calanoida Diaptomidae Skistodiaptomus oregonensis |
| **AY339161** | Calanoida Diaptomidae Skistodiaptomus pygmaeus |
| **MG661001** | Calanoida Eucalanidae Eucalanus bungii |
| **GU969202** | Calanoida Eucalanidae Eucalanus elongatus |
| **GU969148** | Calanoida Eucalanidae Pareucalanus attenuatus |
| **GU969190** | Calanoida Eucalanidae Rhincalanus cornutus |
| **GU969191** | Calanoida Eucalanidae Rhincalanus nasutus |
| **GU969168** | Calanoida Eucalanidae Subeucalanus crassus |
| **GU969169** | Calanoida Eucalanidae Subeucalanus mucronatus |
| **GU969143** | Calanoida Eucalanidae Subeucalanus subtenuis |
| **KR048714** | Calanoida Eurytemora pacifica |
| **MG661044** | Calanoida Heterorhabdidae Heterorhabdus norvegicus |
| **AB625959** | Calanoida Heterorhabdidae Heterorhabdus tanneri |
| **AB625968** | Calanoida Heterorhabdidae Heterostylites major |
| **LC107407** | Calanoida Heterorhabdidae Mesorhabdus brevicaudatus |

| | |
|---|---|
| **MG661043** | Calanoida Heterorhabdidae Paraheterorhabdus compactus |
| **MG660995** | Calanoida Lucicutiidae Lucicutia anomala |
| **AB625965** | Calanoida Lucicutiidae Lucicutia ovaliformis |
| **MG660916** | Calanoida Lucicutiidae Lucicutia polaris |
| **MG660912** | Calanoida Lucicutiidae Lucicutia pseudopolaris |
| **GU969203** | Calanoida Mecynoceridae Mecynocera clausi |
| **AB625967** | Calanoida Metridinidae Metridia asymmetrica |
| **GU969176** | Calanoida Metridinidae Metridia gerlachei |
| **GU969183** | Calanoida Metridinidae Pleuromamma abdominalis |
| **GU969185** | Calanoida Metridinidae Pleuromamma antarctica |
| **GU969184** | Calanoida Metridinidae Pleuromamma borealis |
| **GU969186** | Calanoida Metridinidae Pleuromamma xiphias |
| **HM997057** | Calanoida Nullosetigeridae Nullosetigera auctiseta |
| **JQ911936** | Calanoida Paracalanidae Acrocalanus andersoni |
| **JQ911937** | Calanoida Paracalanidae Acrocalanus gibber |
| **JQ911938** | Calanoida Paracalanidae Acrocalanus gracilis |
| **JQ911939** | Calanoida Paracalanidae Acrocalanus longicornis |
| **GU969201** | Calanoida Paracalanidae Acrocalanus monachus |
| **JQ911941** | Calanoida Paracalanidae Bestiolina similis |
| **GU969180** | Calanoida Paracalanidae Paracalanus aculeatus |
| **JQ911956** | Calanoida Paracalanidae Paracalanus denudatus |
| **JF326205** | Calanoida Paracalanidae Paracalanus parvus |
| **JQ911957** | Calanoida Paracalanidae Paracalanus quasimodo |
| **JQ911958** | Calanoida Paracalanidae Paracalanus tropicus |
| **MG660954** | Calanoida Phaennidae Onchocalanus cristogerens |
| **MF959837** | Calanoida Phaennidae Xanthocalanus LM24 |
| **MG661040** | Calanoida Phaennidae Xanthocalanus polarsternae |
| **MG661041** | Calanoida Phaennidae Xanthocalanus profundus |
| **JX984662** | Calanoida Pseudodiaptomidae Pseudodiaptomus annandalei |
| **JX984663** | Calanoida Pseudodiaptomidae Pseudodiaptomus aurivillii |
| **KC815329** | Calanoida Pseudodiaptomidae Pseudodiaptomus euryhalinus |
| **GU969194** | Calanoida Pseudodiaptomidae Pseudodiaptomus inopinus |
| **KR048712** | Calanoida Pseudodiaptomidae Pseudodiaptomus marinus |
| **GU969193** | Calanoida Pseudodiaptomidae Pseudodiaptomus poplesia |
| **MF959838** | Calanoida Rostrocalanidae Rostrocalanus cognatus |
| **MF959839** | Calanoida Rostrocalanidae Rostrocalanus JR11 |

| | |
|---|---|
| **KU247619** | Calanoida Spinocalanidae Mimocalanus crassus |
| **KU247638** | Calanoida Spinocalanidae Monacilla tenera |
| **KU247634** | Calanoida Spinocalanidae Mospicalanus schielae |
| **MF796501** | Calanoida Spinocalanidae Spinocalanus abyssalis |
| **MG660997** | Calanoida Spinocalanidae Spinocalanus antarcticus |
| **AY192563** | Calanoida Subeucalanus pileatus |
| **KX400986** | Calanoida Temoridae Eurytemora affinis |
| **KX400986** | Calanoida Temoridae Eurytemora affinis |
| **MN567280** | Calanoida Temoridae Eurytemora americana |
| **KX400977** | Calanoida Temoridae Eurytemora carolleeae |
| **GU969209** | Calanoida Temoridae Temora discaudata |
| **JX995310** | Calanoida Temoridae Temora longicornis |
| **GU969210** | Calanoida Temoridae Temora stylifera |
| **GU969211** | Calanoida Temoridae Temora turbinata |
| **HM997066** | Calanoida Temoridae Temoropia mayumbaensis |
| **AB292213** | Chelicerata Pycnogonida Pycnogonum tenue |
| **Hexapoda** | Collembola Crossodonthina koreana |
| **Hexapoda** | Collembola Cyphoderus javanus |
| **Hexapoda** | Collembola Hypogastrura dolsana |
| **Hexapoda** | Collembola Lepidocyrtus paradoxus |
| **Hexapoda** | Collembola Podura aquatica |
| **KR048724** | Cyclopoida Bonnierilla curvicaudata |
| **FJ214952** | Cyclopoida Cyclopettidae Paracyclopina nana |
| **MF077729** | Cyclopoida Cyclopicina longifurcata |
| **AY626999** | Cyclopoida Cyclopidae Acanthocyclops viridis |
| **KR048733** | Cyclopoida Cyclopidae Apocyclops borneoensis |
| **AY626997** | Cyclopoida Cyclopidae Apocyclops royi |
| **EF532821** | Cyclopoida Cyclopidae Cyclops insignis |
| **EF532820** | Cyclopoida Cyclopidae Cyclops kolensis |
| **AJ746335** | Cyclopoida Cyclopidae Eucyclops dumonti |
| **L81940** | Cyclopoida Cyclopidae Eucyclops serrulatus |
| **AJ746333** | Cyclopoida Cyclopidae Eucyclops speratus |
| **KR048727** | Cyclopoida Cyclopidae Megacyclops viridis |
| **MH571998** | Cyclopoida Cyclopidae Mesocyclops pehpeiensis |
| **DQ107580** | Cyclopoida Cyclopidae Thermocyclops WEN |
| **KR048729** | Cyclopoida Cyclopidae Tropocyclops ishidai |

| | |
|---|---|
| **JF288757** | Cyclopoida Oithonidae Oithona brevicornis |
| **KJ814022** | Cyclopoida Oithonidae Oithona davisae |
| **GU969179** | Cyclopoida Oithonidae Oithona similis |
| **KR048731** | Cyclopoida Pachypygus curvatus |
| **KR048766** | Cyclopoida Rhynchomolgidae Critomolgus vicinus |
| **JF781541** | Cyclopoida Rhynchomolgidae Doridicola agilis |
| **KR048761** | Cyclopoida Rhynchomolgidae Zamolgus cavernularius |
| **MN536876** | Harpacticoida Aegisthidae Cerviniella brodskayae |
| **MF077767** | Harpacticoida Aegisthidae Cerviniopsis longicaudata |
| **MN536817** | Harpacticoida Aegisthidae Pontostratiotes fontani |
| **MF077716** | Harpacticoida Ectinosomatidae Bradya DZMB037 |
| **AY627016** | Harpacticoida Ectinosomatidae Bradya Greenland-RJH-2004 |
| **MF077759** | Harpacticoida Ectinosomatidae Parabradya dilatata |
| **EU370430** | Harpacticoida Harpacticidae Tigriopus fulvus |
| **EU054307** | Harpacticoida Harpacticidae Tigriopus japonicus |
| **MK075972** | Harpacticoida Nannopodidae Nannopus bulbiseta |
| **MK075968** | Harpacticoida Nannopodidae Nannopus minutus |
| **MK075984** | Harpacticoida Nannopodidae Nannopus parvipilis |
| **MK075964** | Harpacticoida Nannopodidae Nannopus serratus |
| **MF077760** | Harpacticoida Pseudotachidiidae Pseudotachidius bipartitus |
| **MF077748** | Harpacticoida Pseudotachidiidae Xylora bathyalis |
| **EU368600** | Hexapoda Diplura Dicellurata Catajapyx aquilonaris |
| **AY037168** | Hexapoda Diplura Dicellurata Parajapyx emeryanus |
| **AF173234** | Hexapoda Diplura Rhabdura Campodea tillyardi |
| **AY145134** | Hexapoda Diplura Rhabdura Octostigma sinensis |
| **AY145137** | Hexapoda Diplura Rhabdura Pseudlibanocampa sinensis |
| **EU368613** | Hexapoda Insecta Archaeognatha Machilidae Lepismachilis y-signata |
| **EU368614** | Hexapoda Insecta Archaeognatha Machilidae Pedetontus okajimae |
| **AF370788** | Hexapoda Insecta Archaeognatha Meinertellidae Allomachilis froggatti |
| **JQ581037** | Hexapoda Insecta Coleoptera Diabrotica virgifera |
| **MH746297** | Hexapoda Insecta Coleoptera Hoplopactus lateralis |
| **AY121133** | Hexapoda Insecta Dermaptera Chelisoches morio |
| **KX069007** | Hexapoda Insecta Dermaptera Irdex papuanus |
| **GAYK02035584** | Hexapoda Insecta Mecoptera Boreus hyemalis |
| **AY521864** | Hexapoda Insecta Megaloptera Sialis hamata |
| **JQ581037** | Hexapoda Insecta Neoptera Coleoptera Diabrotica virgifera |

| | |
|---|---|
| **MH746297** | Hexapoda Insecta Neoptera Coleoptera Hoplopactus lateralis |
| **AY121133** | Hexapoda Insecta Neoptera Dermaptera Chelisoches morio |
| **KX069007** | Hexapoda Insecta Neoptera Dermaptera Irdex papuanus |
| **AF440198** | Hexapoda Insecta Neoptera Diptera Anopheles maculatus |
| **AF322422** | Hexapoda Insecta Neoptera Diptera Oestromyia leporina |
| **AF322421** | Hexapoda Insecta Neoptera Diptera Ornithomya avicularia |
| **AF322419** | Hexapoda Insecta Neoptera Diptera Sarcophaga bullata |
| **KT188467** | Hexapoda Insecta Neoptera Hemiptera Bannacoris arboreus |
| **KT188471** | Hexapoda Insecta Neoptera Hemiptera Lestonia haustorifera |
| **KT188473** | Hexapoda Insecta Neoptera Hemiptera Sagriva vittata |
| **AF286289** | Hexapoda Insecta Neoptera Holometabola Mecoptera Chorista Chorista australis |
| **KR068932** | Hexapoda Insecta Neoptera Lepidoptera Aidos perfusa admiranda |
| **MG100879** | Hexapoda Insecta Neoptera Lepidoptera Leguminivora glycinivorella |
| **KR068953** | Hexapoda Insecta Neoptera Lepidoptera Perola murina |
| **AF286296** | Hexapoda Insecta Neoptera Mecoptera Brachypanorpa carolinensis |
| **X89493** | Hexapoda Insecta Neoptera Mecoptera Panorpa germanica |
| **AY521864** | Hexapoda Insecta Neoptera Megaloptera Sialis hamata |
| **Z97565** | Hexapoda Insecta Neoptera Orthoptera Caelifera Cylindraustralia kochii |
| **KM853181** | Hexapoda Insecta Neoptera Orthoptera Caelifera Ellipes minuta |
| **Z97567** | Hexapoda Insecta Neoptera Orthoptera Caelifera Euschmidtia cruciformis |
| **Z97586** | Hexapoda Insecta Neoptera Orthoptera Caelifera Tanaocerus koebeli |
| **Z97566** | Hexapoda Insecta Neoptera Orthoptera Ensidera Cyphoderris monstrosa |
| **Z97564** | Hexapoda Insecta Neoptera Orthoptera Ensifera Comicus campestris |
| **ET713455** | Hexapoda Insecta Neoptera Orthoptera Ensifera Hemiandrus subantarcticus |
| **ET676721** | Hexapoda Insecta Neoptera Orthoptera Ensifera Motuweta riparia |
| **EF622722** | Hexapoda Insecta Neoptera Plecoptera Despaxia augusta |
| **AY121148** | Hexapoda Insecta Neoptera Plecoptera Isoperla davisi |
| **U68400** | Hexapoda Insecta Neoptera Plecoptera Mesoperlina pecircai |
| **Z97595** | Hexapoda Insecta Neoptera Plecoptera Nemoura meieri |
| **EF622698** | Hexapoda Insecta Neoptera Polyneoptera Plecoptera Neuroperla schedingi |
| **X89486** | Hexapoda Insecta Neoptera Siphonaptera Archaeopsylla erinacei |
| **EU336056** | Hexapoda Insecta Neoptera Siphonaptera Rhadinopsylla masculana |
| **AF423886** | Hexapoda Insecta Neoptera Siphonaptera Thrassis bacchi gladiolus |
| **EU336105** | Hexapoda Insecta Neoptera Siphonaptera Uropsylla tasmanica |
| **KM463948** | Hexapoda Insecta Neoptera Trichoptera Asynarchus nigriculus |
| **KM463952** | Hexapoda Insecta Neoptera Trichoptera Limnephilus picturatus |

| | |
|---|---|
| **AF286300** | Hexapoda Insecta Neoptera Trichoptera Oecetis avara |
| **AF286292** | Hexapoda Insecta Neoptera Trichoptera Pycnopsyche lepida |
| **AY521892** | Hexapoda Insecta Neoptera Zoraptera Zorotypus hubbardi |
| **KM246626** | Hexapoda Insecta Neoptera Zoraptera Zorotypus medoensis |
| **AY521891** | Hexapoda Insecta Neoptera Zoraptera Zorotypus novobrittanicus |
| **ET713455** | Hexapoda Insecta Orthoptera Hemiandrus subantarcticus |
| **ET676721** | Hexapoda Insecta Orthoptera Motuweta riparia |
| **AY121148** | Hexapoda Insecta Plecoptera Isoperla davisi |
| **EU368597** | Hexapoda Protura Acerentomata Acerentomon franzi |
| **AY037169** | Hexapoda Protura Acerentomata Baculentulus tienmushanensis |
| **EU557242** | Hexapoda Protura Acerentomata Hesperentomon pectigastrulum |
| **EU557254** | Hexapoda Protura Eosentomata Eosentomon megaglenum |
| **EU557253** | Hexapoda Protura Eosentomata Eosentomon orientale |
| **EU557252** | Hexapoda Protura Eosentomata Zhongguohentomon piligeroum |
| **AY596359** | Hexapoda Protura Sinentomata Fujientomon dicestum |
| **AY596358** | Hexapoda Protura Sinentomata Sinentomon erythranum |
| **EF582897** | Malacostraca Amphipoda Crangonyx pseudogracilis |
| **EF582931** | Malacostraca Amphipoda Gammarus takesensis |
| **AB295408** | Malacostraca Amphipoda Jassa slatteryi |
| **U33181** | Malacostraca Decapoda Astacus astacus |
| **DQ925828** | Malacostraca Decapoda Cymonomoides delli |
| **AY743956** | Malacostraca Decapoda Hippolyte pleuracanthus |
| **MG677849** | Malacostraca Euphausiacea Stylocheiron maximum |
| **MG677856** | Malacostraca Euphausiacea Thysanopoda aequalis |
| **HM138863** | Malacostraca Hoplocarida Coronis scolopendra |
| **KM074038** | Malacostraca Hoplocarida Haptosquilla hamifera |
| **EF116546** | Malacostraca Isopoda Betamorpha fusiformis |
| **KY951738** | Malacostraca Isopoda Ketosoma werneri |
| **EF682249** | Malacostraca Isopoda Notopais magnifica |
| **KY951740** | Malacostraca Isopoda Thaumastosoma platycarpus |
| **MF077710** | Misophrioida Archimisophria discoveryi |
| **MF077709** | Misophrioida Benthomisophria cornuta |
| **MF077778** | Misophrioida Benthomisophria palliata |
| **MF077772** | Misophrioida Misophriella DZMB392 |
| **LC320120** | Misophrioida Misophriidae HT-2017 |
| **JF781532** | Misophrioida Misophriopsis okinawensis |

| | |
|---|---|
| **KR048782** | Monstrilloida Cymbasoma reticulatum |
| **KR048787** | Monstrilloida Maemonstrilla simplex |
| **DQ538495** | Monstrilloida Monstrilla clavata |
| **MF077738** | Monstrilloida Monstrilla helgolandica |
| **KR048783** | Monstrilloida Monstrillopsis SYB-2016 |
| **MF077786** | Mormonilloida Mormonilla DZMBK37 |
| **MF077740** | Mormonilloida Mormonilla phasma |
| **MF077739** | Mormonilloida Neomormonilla minor |
| **KR048768** | Poecilostomatoida Bomolochidae Bomolochus bellones |
| **KR048747** | Poecilostomatoida Bomolochidae Nothobomolochus thambus |
| **JF781555** | Poecilostomatoida Catiniidae Catinia plana |
| **KR048753** | Poecilostomatoida Chondracanthidae Acanthochondria spirigera |
| **L34046** | Poecilostomatoida Chondracanthidae Chondracanthus lophii |
| **MF077732** | Poecilostomatoida Chondracanthidae Chondracanthus merluccii |
| **JF781553** | Poecilostomatoida Clausidiidae Clausidium vancouverense |
| **KR048764** | Poecilostomatoida Clausidiidae Conchyliurus dispar |
| **KR048763** | Poecilostomatoida Clausidiidae Conchyliurus quintus |
| **KR048744** | Poecilostomatoida Clausidiidae Hemicyclops ctenidis |
| **KR048769** | Poecilostomatoida Clausidiidae Hemicyclops tanakai |
| **JF781552** | Poecilostomatoida Clausidiidae Hemicyclops thalassius |
| **GU969165** | Poecilostomatoida Corycaeidae Corycaeus speciosus |
| **MF077730** | Poecilostomatoida Corycaeidae Ditrichocorycaeus anglicus |
| **DQ107577** | Poecilostomatoida Ergasilidae Ergasilus peregrinus |
| **DQ107569** | Poecilostomatoida Ergasilidae Ergasilus tumidus |
| **DQ107578** | Poecilostomatoida Ergasilidae Ergasilus yaluzangbus |
| **DQ107576** | Poecilostomatoida Ergasilidae Paraergasilus brevidigitus |
| **DQ107563** | Poecilostomatoida Ergasilidae Sinergasilus polycolpus |
| **DQ107561** | Poecilostomatoida Ergasilidae Sinergasilus undulatus |
| **MK370210** | Poecilostomatoida Oncaeidae Oncaea 776DZMB |
| **MG661013** | Poecilostomatoida Oncaeidae Oncaea notopus |
| **MG661033** | Poecilostomatoida Oncaeidae Triconia borealis |
| **GU969173** | Poecilostomatoida Sapphirinidae Sapphirina darwinii |
| **GU969208** | Poecilostomatoida Sapphirinidae Sapphirina scarlata |
| **MF077781** | Progymnoplea Platycopioida Platycopia perplexa |
| **MF077704** | Progymnoplea Platycopioida Nanocopia minuta |
| **FJ447452** | Siphonostomatoida Achtheinus oblongus |

| | |
|---|---|
| **EF088411** | Siphonostomatoida Caligus pelamydis |
| **KR048772** | Siphonostomatoida Hatschekia japonica |
| **EF088414** | Siphonostomatoida Lepeophtheirus pollachius |
| **FJ447428** | Siphonostomatoida Nemesis lamna |
| **FJ447445** | Siphonostomatoida Nesippus orientalis |
| **LC054034** | Siphonostomatoida Rhizorhina ohtsukai |

**Table S7.** Reference sequences used in the phylogenetic tree of tunicates (Fig. S7).

| Accession code | Simplified taxonomic lineage |
|---|---|
| **MK621850** | Tunicata Appendicularia Copelata Oikopleuridae Oikopleura_gorskyi |
| **MK621851** | Tunicata Appendicularia Copelata Oikopleuridae Oikopleura_cophocerca |
| **MK621864** | Tunicata Appendicularia Copelata Oikopleuridae Mesoikopleura_haranti |
| **MK621845** | Tunicata Appendicularia Copelata Oikopleuridae Oikopleura_parva |
| **MK621852** | Tunicata Appendicularia Copelata Oikopleuridae Oikopleura_labradoriensis |
| **MG661055** | Tunicata Appendicularia Copelata Oikopleuridae Oikopleura_vanhoeffeni |
| **MK621854** | Tunicata Appendicularia Copelata Oikopleuridae Folia_mediterranea |
| **L12434** | Tunicata Ascidiacea Stolidobranchia Molgulidae Molgula Molgula_provisionalis |
| **L12430** | Tunicata Ascidiacea Stolidobranchia Molgulidae Molgula Molgula_occulta |
| **L12432** | Tunicata Ascidiacea Stolidobranchia Molgulidae Molgula Molgula_oculata |
| **L12418** | Tunicata Ascidiacea Stolidobranchia Molgulidae Molgula Molgula_bleizi |
| **L12441** | Tunicata Ascidiacea Stolidobranchia Styelidae Polycarpa Polycarpa_pomaria |
| **L12413** | Tunicata Ascidiacea Stolidobranchia Styelidae Cnemidocarpa Cnemidocarpa_finmarkiensis |
| **L12422** | Tunicata Ascidiacea Stolidobranchia Molgulidae Molgula Molgula_complanata |
| **L12420** | Tunicata Ascidiacea Stolidobranchia Molgulidae Molgula Molgula_citrina |
| **AB013013** | Tunicata Thaliacea Doliolida Doliolidae Doliolum Doliolum_nationalis |
| **AB013011** | Tunicata Thaliacea Pyrosomata Pyrosomatidae Pyrosoma Pyrosoma_atlanticum |
| **AB013012** | Tunicata Thaliacea Doliolida Doliolidae Doliolum Doliolum_nationalis |
| **AB013015** | Tunicata Appendicularia Copelata Oikopleuridae Oikopleura Oikopleura_sp. |
| **AB013016** | Tunicata Ascidiacea Stolidobranchia Pyuridae Halocynthia Halocynthia_roretzi |
| **X53538** | Tunicata Ascidiacea Stolidobranchia Pyuridae Herdmania Herdmania_momus |
| **AB211066** | Tunicata Ascidiacea Stolidobranchia Styelidae Botryllus Botryllus_schlosseri |
| **JQ917223** | Tunicata Thaliacea Salpida Salpidae Pegea Pegea_confoederata |
| **KT387603** | Tunicata Ascidiacea Stolidobranchia Pyuridae Microcosmus Microcosmus_exasperatus |
| **FM244859** | Tunicata Ascidiacea Stolidobranchia Styelidae Asterocarpa Asterocarpa_humilis |
| **FM244856** | Tunicata Ascidiacea Stolidobranchia Pyuridae Pyura Pyura_dura |
| **FM244865** | Tunicata Thaliacea Salpida Salpidae Ihlea Ihlea_racovitzai |
| **FM244854** | Tunicata Ascidiacea Stolidobranchia Pyuridae Microcosmus Microcosmus_sabatieri |
| **FM244863** | Tunicata Thaliacea Pyrosomata Pyrosomatidae Pyrosomella Pyrosomella_verticillata |
| **FM244855** | Tunicata Ascidiacea Stolidobranchia Pyuridae Microcosmus Microcosmus_squamiger |
| **FM244869** | Tunicata Appendicularia Copelata Oikopleuridae Oikopleura Oikopleura_labradoriensis |
| **FM244850** | Tunicata Ascidiacea Stolidobranchia Molgulidae Molgula Molgula_occidentalis |
| **FM244868** | Tunicata Appendicularia Copelata Oikopleuridae Megalocercus Megalocercus_huxleyi |

| | |
|---|---|
| **FM244857** | Tunicata Ascidiacea Stolidobranchia Pyuridae Pyura Pyura_gangelion |
| **FM244851** | Tunicata Ascidiacea Stolidobranchia Pyuridae Halocynthia Halocynthia_spinosa |
| **FM244860** | Tunicata Ascidiacea Stolidobranchia Styelidae Polycarpa Polycarpa_mytiligera |
| **KT881545** | Tunicata Appendicularia Copelata Oikopleuridae Bathochordaeus Bathochordaeus_charon |
| **JF961807** | Tunicata Ascidiacea Stolidobranchia Pyuridae Pyura Pyura_praeputialis |
| **FM244853** | Tunicata Ascidiacea Stolidobranchia Pyuridae Microcosmus Microcosmus_polymorphus |
| **JF961794** | Tunicata Ascidiacea Stolidobranchia Pyuridae Pyura Pyura_herdmani |
| **KR057223** | Tunicata Thaliacea Salpida Salpidae Brooksia Brooksia_lacromae |
| **JN565044** | Tunicata Ascidiacea Stolidobranchia Hexacrobylidae Oligotrema Oligotrema_lyra |
| **AY903922** | Tunicata Ascidiacea Stolidobranchia Styelidae Metandrocarpa Metandrocarpa_taylori |
| **AY903926** | Tunicata Ascidiacea Stolidobranchia Pyuridae Pyura Pyura_haustor |
| **AY903923** | Tunicata Ascidiacea Stolidobranchia Styelidae Styela Styela_gibbsii |
| **AY903925** | Tunicata Ascidiacea Stolidobranchia Pyuridae Halocynthia Halocynthia_igaboja |
| **AY903924** | Tunicata Ascidiacea Stolidobranchia Pyuridae Boltenia Boltenia_villosa |
| **AY903921** | Tunicata Ascidiacea Stolidobranchia Molgulidae Molgula Molgula_retortiformis |
| **AY903927** | Tunicata Ascidiacea Stolidobranchia Styelidae Botrylloides Botrylloides_violaceus |
| **AF165827** | Tunicata Ascidiacea Stolidobranchia Pyuridae Herdmania Herdmania_momus |
| **HQ015387** | Tunicata Thaliacea Salpida Salpidae Pegea Pegea_confoederata |
| **HQ015406** | Tunicata Thaliacea Salpida Salpidae Salpa Salpa_thompsoni |
| **HQ015403** | Tunicata Thaliacea Salpida Salpidae Brooksia Brooksia_rostrata |
| **HQ015389** | Tunicata Thaliacea Salpida Salpidae Soestia Soestia_zonaria |
| **HQ015411** | Tunicata Thaliacea Salpida Salpidae Ritteriella Ritteriella_retracta |
| **HQ015386** | Tunicata Thaliacea Salpida Salpidae Pegea Pegea_confoederata |
| **HQ015405** | Tunicata Thaliacea Salpida Salpidae Salpa Salpa_aspera |
| **KJ818250** | Tunicata Ascidiacea Stolidobranchia Styelidae Styela Styela_plicata |
| **HQ015394** | Tunicata Thaliacea Salpida Salpidae Cyclosalpa Cyclosalpa_polae |
| **HQ015415** | Tunicata Thaliacea Salpida Salpidae Thalia Thalia_democratica |
| **HQ015390** | Tunicata Thaliacea Salpida Salpidae Thetys Thetys_vagina |
| **HQ015379** | Tunicata Thaliacea Pyrosomata Pyrosomatidae Pyrostremma Pyrostremma_spinosum |
| **HQ015400** | Tunicata Thaliacea Salpida Salpidae Iasis Iasis_cylindrica |
| **DQ346655** | Tunicata Ascidiacea Stolidobranchia Styelidae Symplegma Symplegma_viride |
| **D14366** | Tunicata Thaliacea Salpida Salpidae Thalia Thalia_democratica |
| **DQ346654** | Tunicata Ascidiacea Stolidobranchia Styelidae Polycarpa Polycarpa_papillata |
